# Geography is more important than life history in the recent diversification of the tiger salamander complex

**DOI:** 10.1101/2020.07.10.197624

**Authors:** Kathryn M. Everson, Levi N. Gray, Angela G. Jones, Nicolette M. Lawrence, Mary E. Foley, Kelly L. Sovacool, Justin D. Kratovil, Scott Hotaling, Paul M. Hime, Andrew Storfer, Gabriela Parra-Olea, Ruth Percino-Daniel, X. Aguilar-Miguel, Eric M. O’Neill, Luis Zambrano, H. Bradley Shaffer, David W. Weisrock

## Abstract

The North American tiger salamander species complex, including its best-known species, the Mexican axolotl, has long been a source of biological fascination. The complex exhibits a wide range of variation in developmental life history strategies, including populations and individuals that undergo metamorphosis, those able to forego metamorphosis and retain a larval, aquatic lifestyle (i.e., paedomorphosis), and those that do both. This life history variation has been assumed to lead to reproductive isolation and speciation, but the degree to which it has shaped population- and species-level divergence is poorly understood. Using a large multi-locus dataset from hundreds of samples across North America, we identified genetic clusters across the geographic range of the tiger salamander complex. These clusters often contain a mixture of paedomorphic and metamorphic taxa, indicating that geographic isolation has played a larger role in lineage divergence than paedomorphosis in this system. This conclusion is bolstered by geography-informed analyses indicating no effect of life history strategy on population genetic differentiation and by model-based analyses demonstrating gene flow between adjacent metamorphic and paedomorphic populations. This fine-scale genetic perspective on life-history variation establishes a framework for understanding how plasticity, local adaptation, and gene flow contribute to lineage divergence. Many members of the tiger salamander complex are endangered, and the Mexican axolotl is an important model system in regenerative and biomedical research. Our results chart a course for more informed use of these taxa in experimental, ecological, and conservation research.

**Significance Statement:** Population structure and speciation are shaped by a combination of biotic and abiotic factors. The tiger salamander complex has been considered to be a key group where life history variation has led to a rapid rate of speciation, driven in large part by the evolution of obligate paedomorphosis–a condition where adults maintain an aquatic, larval phenotype. Using a large multi-locus dataset, we present evidence of gene flow between taxa with different life history strategies, suggesting that obligate paedomorphosis is not a strong driver of speciation in the tiger salamander complex. Many of these nominal taxa are listed as critically endangered, and our genetic results provide information and guidance that will be useful for their conservation.

## Introduction

Life history—the complement of traits affecting survival and reproduction over an organism’s lifetime—affects ecology and dispersal, and therefore plays a potentially important role in shaping population structure and speciation (1, 2). The Mexican axolotl (*Ambystoma mexicanum;* hereafter, “axolotl”) and related salamander species of the North American *Ambystoma tigrinum* complex (Table 1) display enormous life history variation, particularly with respect to the completion of metamorphosis (3). Some species are considered obligate paedomorphs, where sexually mature adults retain a larval, aquatic body plan that includes external gills and an enlarged tail fin (4). Others are considered obligate metamorphs, transforming from aquatic larvae to terrestrial juveniles, eventually returning to the water to breed as adults. Most populations are facultative, transforming under certain genetic and/or environmental conditions (5–8). This lability in life history is believed to have played an important role in the diversification of the *A. tigrinum* complex, particularly in the Trans-Mexican Volcanic Belt (TMVB) region, where several paedomorphic species, including the axolotl, are currently recognized (9–11). Obligate paedomorphosis is estimated to have evolved in this region multiple times (11–13) in association with relatively large, permanent bodies of water. Presumably, restricted gene flow between these isolated, paedomorphic populations led to speciation as well as morphological adaptations to the aquatic lifestyle (14).

**Table 1.** Currently recognized species in the *Ambystoma tigrinum* complex. Habitat preference, range, conservation status (“Status”), and life history strategy were compiled from AmphibiaWeb (2019) and the IUCN (2019). Conservation status abbreviations: LC = least concern, V = vulnerable, E = endangered, CE = critically endangered, DD = data deficient. Life history codes: 1 = obligate metamorph (no paedomorphs documented in the field), 2 = Strong bias towards metamorphosis with rare paedomorphs found in the field, 3 = Both metamorphosis and paedomorphosis common in the field, 4 = Strong bias towards paedomorphosis with rare metamorphs found in the field, 5 = obligate paedomorph (no metamorphs documented in the field). In the text, “paedomorph” describes taxa scored as 4 or 5, while “facultative paedomorph” describes taxa scored as 2 or 3.

| Species | Habitat Preference | Range | Status | Life History |
| --- | --- | --- | --- | --- |
| *Ambystoma altamirani* Dugès, 1895 | Streams, occasional ponds | High elevations in the west and south of the Valley of Mexico (Mexico State, Morelos, and Distrito Federal) | E | 2 |
| *Ambystoma amblycephalum* Taylor, 1940 | Ponds | Small area of Tacicuario (north and west Michoacan) | CE | 2 |
| *Ambystoma andersoni* Krebs & Brandon, 1984 | Lakes / Ponds | Restricted to Lake Zacapu and its outflow streams (north-central Michoacan) | CE | 5 |
| *Ambystoma bombypellum* Taylor, 1940 | Lakes / Ponds, occasional streams | Known only from type locality San Martin/Asunción, 15 km west of Morelia (Michoacan) | DD | 2 |
| *Ambystoma californiense* Gray 1853 | Vernal pools | Great Central Valley and adjacent inner coast ranges, central California | V | 1 |
| *Ambystoma dumerilii* (Dugès, 1870) | Lakes / Ponds | Restricted to Lake Patzcauro (north-central Michoacan) | CE | 5 |
| *Ambystoma flavipiperatum* Dixon, 1963 | Lakes / Ponds, occasional streams | Sierra de Quila (Jalisco). Historically known from the type locality in Santa Cruz, southwest of Guadalajara. | E | 3 |
| *Ambystoma granulosum* Taylor, 1944 | Lakes / Ponds, occasional streams | Northwest of Toluca city (Mexico State and border of Michoacan) | CE | 3 |
| *Ambystoma leorae* (Taylor, 1943) | Streams | High elevations of Rio Frio in Iztaccihuatl -Popocatepetl Natl. Park (Mexico State and adjacent Puebla) | CE | 2 |
| *Ambystoma lermaense* (Taylor, 1940) | Lakes / Ponds | Highlands near Toluca, in Rio Lerma and Lake Lerma near Almoloya (Mexico State) | E | 3 |
| *Ambystoma mavortium* Baird, 1850* | Lakes / Ponds | Widespread distribution from west-central Canada through the USA to northern Mexico (Sonora, Chihuahua, Coahuila). Disjunct distribution in northwestern USA and Canada (Washington, British Columbia). | LC | 3 |
| *Ambystoma mexicanum* (Shaw and Nodder, 1798) | Lakes | Southern areas of Xochimilco (Mexico City). Historically distributed in Lakes of Xochimilco and Chalco. | CE | 4 |
| *Ambystoma ordinarium* Taylor, 1940 | Streams | From south of Patzcuaro through Morelia City to eastern Michoacan | E | 3 |
| *Ambystoma rivulare* (Taylor, 1940) | Streams | Sierra Chinchua, Mariposa Monarca Biosphere Reserve, Michoacan, Valle de Bravo, Amanalco and Nevado de Toluca (Mexico State) | E | 2 |
| *Ambystoma rosaceum* Taylor, 1941 | Streams | Sierra Madre Occidental (Sonora, Durango, Chihuahua, Sinaloa, Nayarit, Jalisco) | LC | 3 |
| *Ambystoma silvense* Webb, 2004 | Lakes / Ponds | Sierra Madre Occidental (at least Chihuahua, Durango) | DD | 3 |
| *Ambystoma taylori* Brandon et al., 1982 | Saline lake | Restricted to Lake Alchichica (Puebla) | CE | 4 |
| *Ambystoma tigrinum* Green, 1825 | Lakes / Ponds | From southeast Canada south to east Texas and extending east of the Appalachian Mountains through the Atlantic Coastal Plain north to Long Island (New York) | LC | 2 |
| *Ambystoma velasci* (Dugès, 1888) | Lakes / Ponds, Streams | Sierra Madre Occidental (Chihuahua) and south across central Mexico (Durango, Jalisco, Nuevo León, Coahuila, Zacatecas, San Luis Potosi, Hidalgo, Queretaro, Tlaxcala, Puebla, Veracruz, Aguascalientes) | LC | 3 |

The term “species complex,” while not a formal taxonomic category, is often used to describe groups of closely related lineages early in the divergence process. The tiger salamander species complex has been highlighted as a potentially valuable example of such a recent radiation (15) that could provide insight into the early mechanisms initiating and/or maintaining diversity (16–21). However, species complexes pose a number of challenges: phenotypic differences among lineages may be subtle or absent (i.e., cryptic species), ancestral polymorphisms may lead to incomplete lineage sorting that confounds molecular systematic studies, and reproductive barriers may be incomplete. The reliance on phenotype for species diagnosis in the *A. tigrinum* complex may require particular scrutiny, as several phenotypic traits, including developmental state and adult color pattern, can be plastic and highly variable (22–24). Given these challenges, fundamental questions remain concerning the reality of component lineages within the tiger salamander complex, calling into question the current taxonomy as an accurate reflection of its underlying evolutionary history.

Previous research has produced mixed results regarding levels of genetic differentiation among lineages in the *A. tigrinum* complex, as well as the relative importance of paedomorphosis as a driver of diversification. While population- and species-level structure is evident, an important theme emerging across multiple studies has been that reproductive barriers are porous (11, 25, 26). Shaffer and McKnight (11) noted “the striking lack of differentiation among the 14 species of the tiger salamander complex” (p. 425). Some studies have found an elevated degree of population structure among paedomorphic populations relative to metamorphic populations (12, 27). However, others have shown a lack of effect of paedomorphosis on population genetic differentiation (28), and demonstrated that paedomorphic taxa and neighboring metamorphic populations interbreed (29). Furthermore, crosses between metamorphic and paedomorphic taxa can produce viable hybrid offspring under laboratory conditions (30, 31).

To understand the processes underlying diversification in the tiger salamander complex, an important first step must be a range-wide assessment of population structure and the clarification of population- and species-level boundaries. With this groundwork, insights into the role of paedomorphosis and diversification can be addressed. For a study system of this geographic and taxonomic scale, robust inference of population structure requires information from multiple genes combined with thorough range-wide sampling. To meet these criteria, we expand on a large multi-locus dataset containing 95 nuclear loci for 93 individuals (32), to produce a data matrix for 347 individuals across the full geographic range of tiger salamanders (Fig. S1). Given the complex and somewhat checkered history of the group’s taxonomy, we focus on a naïve approach, performing population structure analyses without *a priori* identification of taxa to resolve geographic genetic clusters and characterize patterns of admixture. We also overlay these results on the existing taxonomy to explore the correspondence between current taxonomy and genetic differentiation. We then use these data for a phylogenetic analysis to provide an updated working hypothesis for the evolution of the group. Finally, we test the hypothesis that life history evolution has driven speciation in the tiger salamander complex, particularly through the assumed isolation of paedomorphic species. In light of the life history variation in this species complex, we use natural history information to place taxa into one of five categories (Table 1) reflecting their propensity to metamorphose, then use model-based tests of migration to ask whether paedomorphic populations (categories 4 and 5) exhibit greater levels of genetic differentiation relative to those that regularly metamorphose (categories 1-3).

## Results and Discussion

### Identification of genetic lineages and phylogenetic relationships

A principal components analysis (PCA) and a discriminant analysis of principal components (DAPC; 33) both recovered *Ambystoma californiense* Gray 1853 as genetically distinct from all other tiger salamanders (Fig. 1A, S2, S3), consistent with previous work (e.g., 11, 34). Given that *A. californiense* is the only obligate metamorph in the complex (life history category 1; Table 1) and is genetically divergent and geographically isolated from all other tiger salamanders evaluated in this study, we focus the remaining analyses on the other members of the complex. After removing *A. californiense* from the dataset, DAPC supported the recognition of three primary genetic clusters: U.S. (including samples from southern Canada), northwestern Mexico (primarily the Sierra Madre Occidental), and the central Mexican highlands (Fig. 1B, S4). A PCA of these data produced similar clustering results, with ordination patterns largely mirroring geographic sampling (Fig. S5). For a further discussion of the taxonomic implications of our results, see the Supplementary Material.

**Figure 1.**
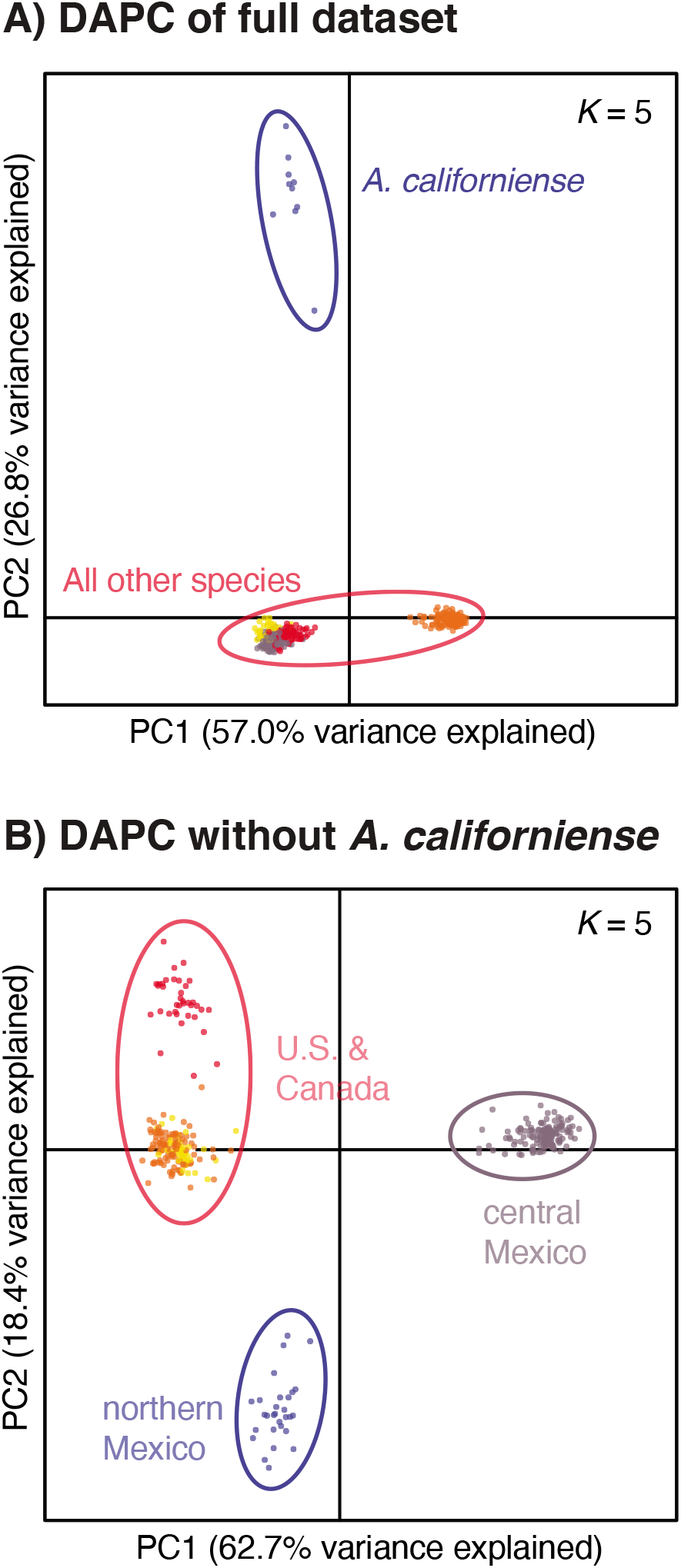
Discriminant analysis of principal components (DAPC) on (A) the entire genomic dataset (without *A. opacum* and *A. texanum* outgroups), and (B) the genomic dataset without *A. californiense*. In both analyses, patterns of genetic differentiation stabilized at values of *K* ≥ 5 (*K* = 5 is shown in both plots, with colors representing genetic clusters). Points represent individuals and ellipses show the groups identified by DAPC. The first and second principal components from the DAPC are on the x and y axes, respectively. Results from additional values of *K* are provided in Supplementary Figures 3 and 4.

Within central Mexico, we identified *K* = 7 as the best-fit number of clusters in both DAPC and STRUCTURE analyses (Figs. 2A, S6). However, several of these clusters were only represented as a small proportion of genetic assignments, rather than as meaningful groups of individuals, and visual inspection at multiple levels of *K* revealed four clear geographic clusters (hereafter referred to as CM1-CM4) that captured the primary patterns of genetic differentiation and admixture across this region (Figs. 2A, S7-S8). Network analysis (35) also recovered groups that generally correspond to CM1-CM4, but with evidence of many reticulations (Fig. 3). This starburst-like pattern indicates the presence of conflicting topologies within the dataset (35), which can be caused by biological processes including recent diversification with incomplete lineage sorting, current or evolutionarily recent gene flow, or both. Phylogenetic analyses (36–38) recovered monophyletic CM1, CM2, and CM4 clusters (see below for discussion of CM3), but often with low statistical support (Fig. 3).

**Figure 2.**
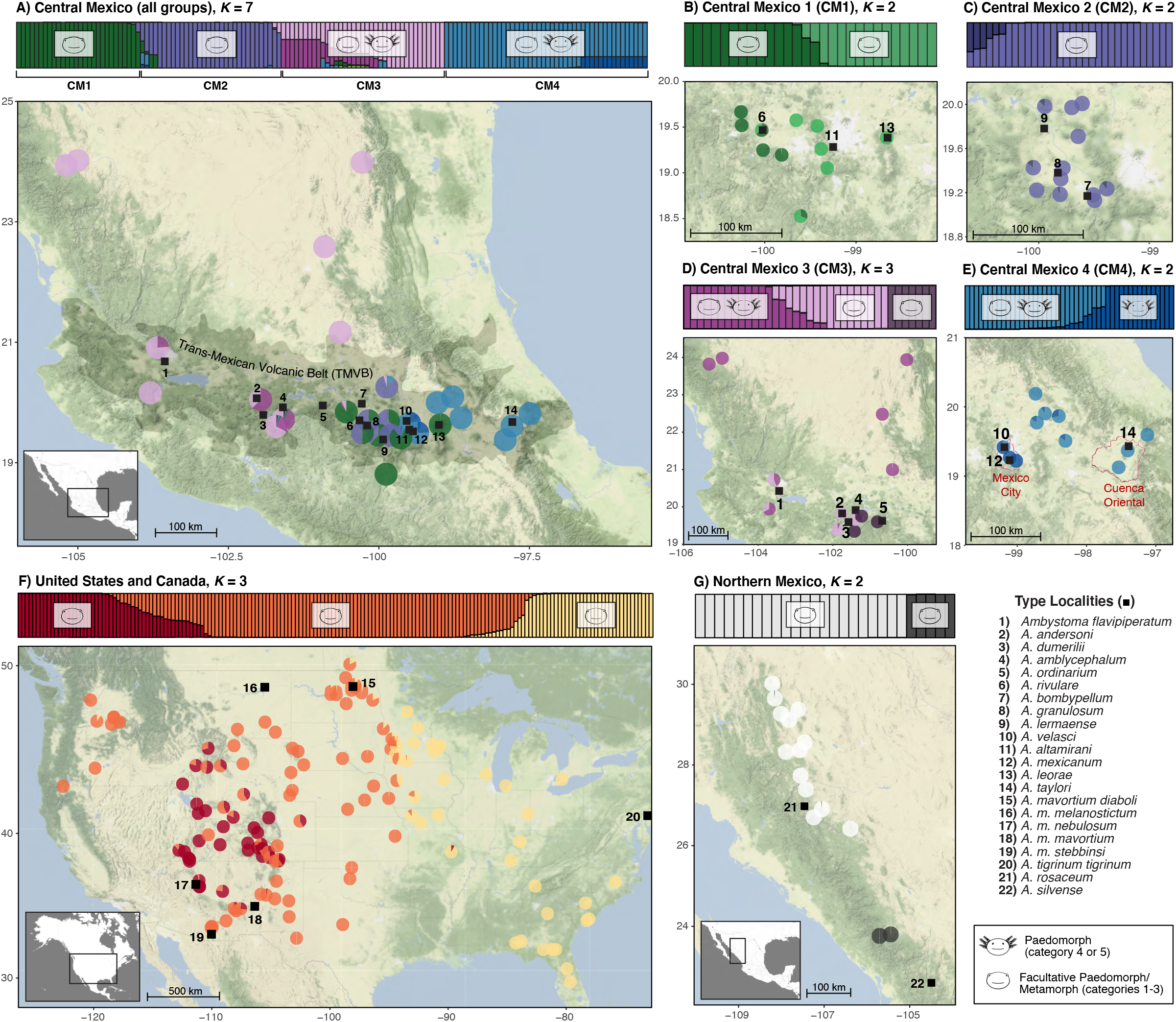
Population genomic analyses of the *Ambystoma tigrinum* species complex. (A) STRUCTURE plot of central Mexico samples. While the Evanno et al. (2005) method identified *K* = 7, four major geographic genetic groups are apparent and were analyzed separately in subsequent analyses (B-E). In each plot, vertical bars represent individuals, while the y-axis shows the membership probability for each group. Pie charts mapped below the assignment plot show the average group membership probability for that locality. Some localities were combined to a single pie chart to reduce visual clutter; see Table S1 for specific coordinates of each sample. The range of the Trans-Mexican Volcanic Belt (TMVB) is identified by shading and is modified from Ferrari (2004). The range of the Cuenca Oriental is outlined in red and is modified from Percino-Daniel et al. (2016). Two geographic outliers placed in CM2 are thought to represent range introductions and are not shown here, but are discussed in supplementary text and shown in Fig. S9. (F,G) STRUCTURE results from analyses of the U.S. and northern Mexico groups, respectively. Detailed results are shown in Figs. S11-S12. Black squares on the maps denote the type localities of all species included in this study. On each STRUCTURE membership plot, salamander face symbols denote the presence of salamanders with a life history category of 1-3 (the cartoon lacks gills) or 4-5 (paedomorphic, cartoon shows gills); groups labeled with both symbols contain a mixture of these life history categories.

**Figure 3.**
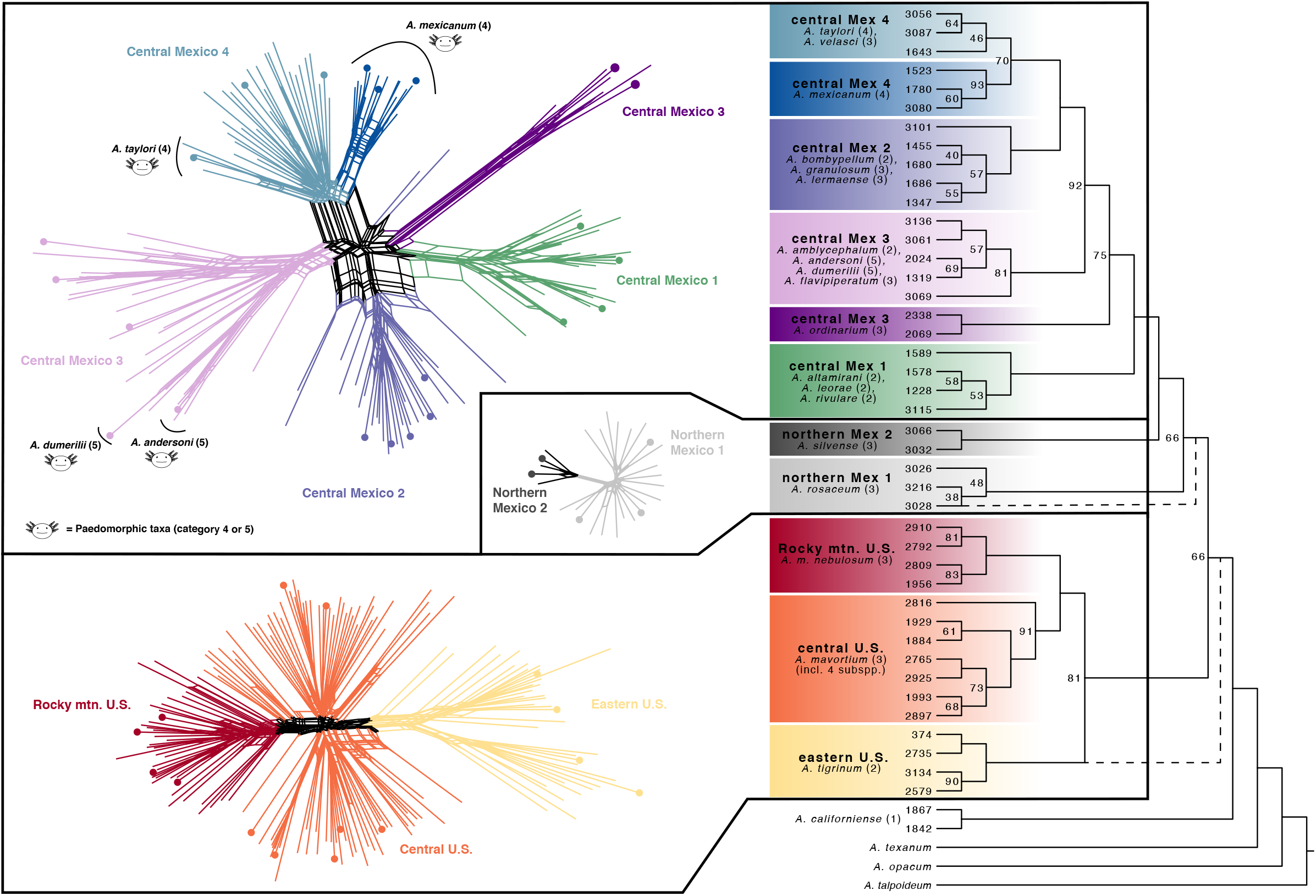
Results of phylogenetic network analyses (left) and the SVDquartets-based species tree analysis (right) of the *Ambystoma tigrinum* species complex. Clades are colored according to their cluster assignments in population genetic analyses, as in Fig. 2. In phylogenetic networks, species names are only shown for paedomorphic taxa in categories 4 or 5 (Table 1). Life history categories for all taxa are given in parentheses in the species tree. Note that SVDquartet analysis used a subset of individuals, indicated by solid circles on the phylogenetic networks. Phylogenetic results from Bayesian and maximum-likelihood concatenated analyses were largely concordant with the quartets tree (Fig. S13), although two important differences are indicated by dashed lines. Bootstrap support (BS) values are indicated on each node of the quartets tree (BS values ≥ 95 are not shown), while tips are labeled with individual IDs (Table S1). Branch lengths are not scaled to time or substitution rate.

Based on type locality and range information, the CM1 cluster includes three stream-breeding and facultatively paedomorphic taxa: *Ambystoma altamirani* Dugès 1895 (39), *A. leorae* (Taylor 1943) (40), and *A. rivulare* (Taylor 1940) (9). Subsequent analyses of this cluster identified two admixed populations associated with eastern and western portions of its geographic distribution (Figs. 2B, S9). Phylogenetically, CM1 was monophyletic and sister to all other central Mexican groups (Fig. 3).

CM2 grouped *A. lermaense* (Taylor 1940) (9) with samples corresponding to the type localities and range of *A. bombypellum* Taylor 1940 (9) and *A. granulosum* Taylor 1944 (41). There was no evidence for genetic isolation among any of these three taxa (Figs. 2C, S9).

Within CM3, population-genetic analyses first separated samples of *A. ordinarium* Taylor 1940 (5) from the rest (with no admixture detected; Figs. 2D, S9). Other samples clustered into northern and southern groups with some admixture in intermediate localities. The more southern cluster includes two obligately paedomorphic (category 5) taxa, *A. andersoni* Krebs & Brandon 1984 (42) and *A. dumerilii* (Dugès 1870) (43), endemic to Lakes Zacapú and Pátzcuaro, respectively. The samples from other CM3 localities likely represent what are currently classified as *A. amblycephalum* Taylor 1940 (9) and *A. flavipiperatum* Dixon 1963 (44).

CM4 included the axolotl, *A. mexicanum* (Shaw & Nodder 1798) (45), and *A. taylori* Brandon et al. 1981 (10)—both category 4 paedomorphs—as well as samples assigned to the facultatively paedomorphic (category 3) *A. velasci* (Duges 1888) (46). Subsequent DAPC and STRUCTURE analyses of CM4 placed all but one *A. mexicanum* individual sampled from Lakes Xochimilco and Chapultepec (i.e., the remaining habitats of the axolotl) in a distinct cluster (Fig. 2E, S9), with evidence of some admixture in samples to the northwest of Mexico City.

Within the U.S. and Canada (excluding *A. californiense),* DAPC and STRUCTURE identified three genetic clusters (*K* = 3), with clear signs of admixed contact zones (Figs. 2F, S10, S11). Geographically, these clusters are associated with the eastern U.S., the central + western U.S., and the southwestern U.S. + Rocky Mountains (hereafter, we refer to the latter two groups as central U.S. and Rocky Mountain U.S., respectively). Populations in the eastern U.S. are assignable to *A. tigrinum* Green 1825, while central U.S. and Rocky Mountain U.S. populations to *A. mavortium* Baird 1850. Phylogenetic analyses corroborated these results, albeit with extensive reticulations among clusters in the phylogenetic network (Fig. 3) and mixed support for species-tree relationships among the three major groups (Fig. 3 vs. Fig. S13). Further exploration of population structure within each of the three U.S. clusters recovered additional patterns of differentiation (Figs. S10, S11), including northern and southern clusters in the eastern U.S. and central U.S., and an allopatric cluster restricted to the Pacific Northwest. Overall, our U.S. results are similar to mtDNA-based results from previous studies, which identified haplotype clades associated with the eastern U.S., Great Plains and Rocky Mountains, and the Pacific Northwest (11, 47).

All analyses of the northern Mexico group identified two genetic clusters (*K* = 2; Figs. 2G, S12). The northern cluster corresponds to *A. rosaceum* Taylor 1941 (48), while the southern cluster is likely *A*. *silvense* Webb 2004 (49, see also 50). Phylogenetic analyses recovered each of these two northern Mexico clusters as monophyletic, but not sister to one another, and with few reticulate nodes in the phylogenetic network (Fig. 3, S13).

### Does paedomorphosis lead to increased genetic differentiation?

As a first step in addressing this question, we used Bayesian modeling to test for genetic isolation of paedomorphic taxa (i.e., species scored as life history category 4 or 5 in Table 1). We tested for gene flow between paedomorphs-*A. andersoni* and *A. dumerilii* in CM3, and *A. mexicanum* and *A. taylori* in CM4-and their surrounding facultatively paedomorphic populations. For both CM3 and CM4, the top-ranking models included migration among all populations, regardless of their degree of paedomorphosis (Fig. 4). Alternative models of no migration or restricted migration (Fig. S14) were rejected with decisive Bayes factors (Table S3). In CM3, migration rates entering the *A. dumerilii* population were higher than those leaving that population, while migration rates entering and exiting *A. andersoni* were approximately equal. In CM4, migration rates exiting the *A. mexicanum* population were much higher than those entering the population while migration rates entering and exiting *A. taylori* were approximately equal (Fig. 4, Table S4). An important caveat to our migration and population structure analyses is that divergence time is not included as a model parameter, which prevents distinguishing between historical and contemporary gene flow. Thus, our ability to differentiate ongoing from historical gene flow in these tests is limited.

**Figure 4.**
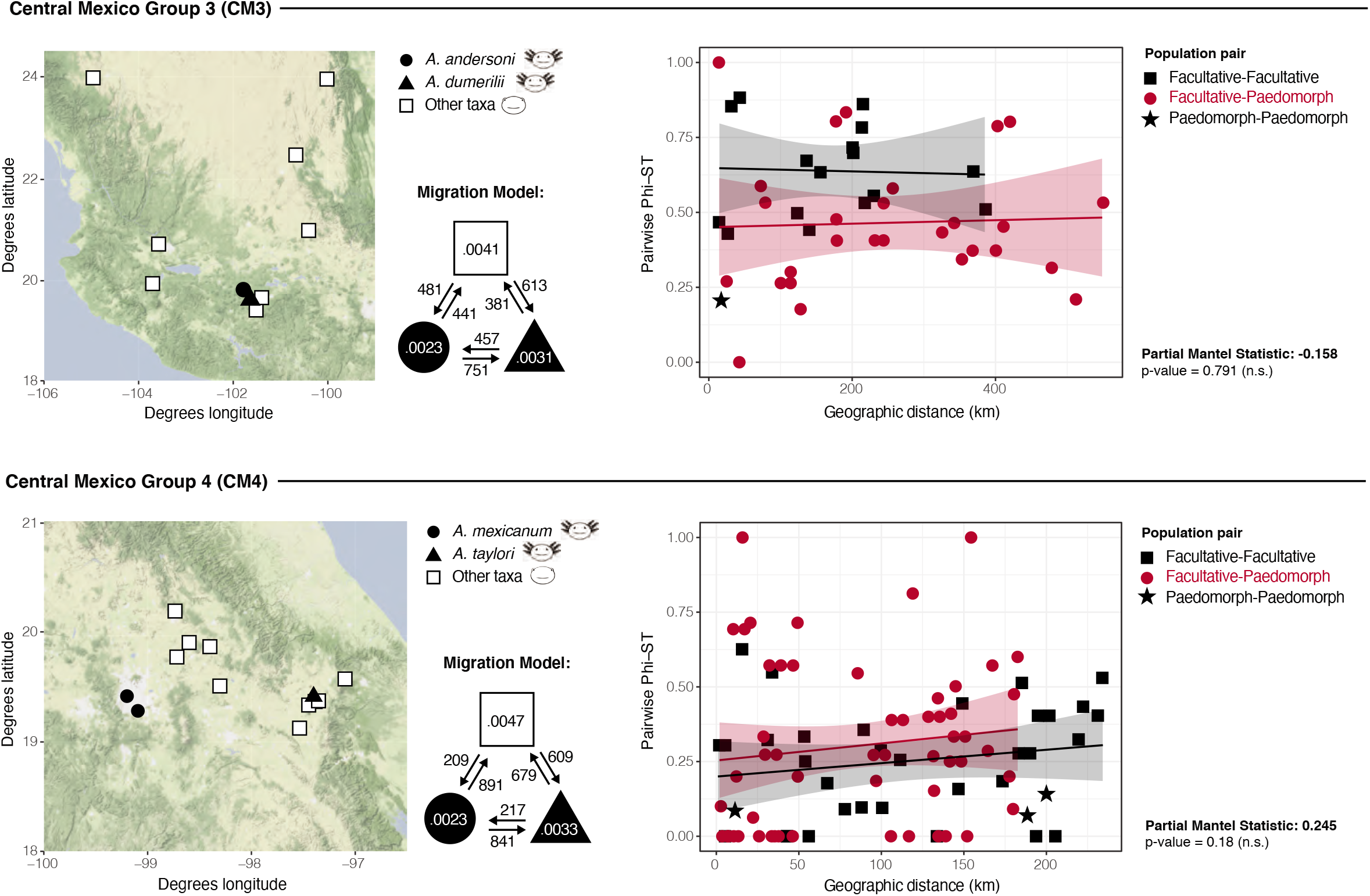
Maps (left) show localities for CM3 and CM4, the two genetic clusters containing paedomorphic taxa (life history categories 4 and 5). The top-ranked migration model is shown to the right of each map, with median mutation-scaled effective population sizes (Θ) shown inside the shapes denoting populations. Arrows indicate the direction of migration (*M* = *m*/μ, where *m* is the fraction of immigrants in each generation and μ is the mutation rate per generation per site) and median migration rates are provided next to each arrow. Full model test results are provided in Table S3 and full parameter estimates are provided in Table S4. Scatterplots (right) show pairwise geographic distances by pairwise Φ_ST_ among all populations. Point colors and shapes denote the life history strategies of the pairwise comparison being made: black squares = both populations are facultatively paedomorphic; red circles = one population is facultatively paedomorphic and one population is obligately paedomorphic; black stars = both populations are obligately paedomorphic. Solid lines denote the line of best fit, calculated separately for each life history comparison. Regressions were not performed on paedomorph-paedomorph comparisons due to low sample sizes. Shaded areas denote the 95% confidence interval for the regression. P-values to the right of each graph were calculated using a partial mantel test, which tested for a correlation between Φ_ST_ and life history while correcting for geographic distance (n.s. = not significant).

We also assessed whether a paedomorphic life history is associated with greater population genetic differentiation (calculated here as Φ_ST_), which would be expected if paedomorphosis is an important reproductive barrier (12). Plots of Φ_ST_ against geographic distance showed that comparisons involving paedomorphic populations were largely overlapping with those based on facultative-facultative comparisons, with no difference in slope or y-intercept (Fig. 4). In both CM3 and CM4, partial Mantel tests found no significant correlation between life history and pairwise Φ_ST_ after correcting for geographic distance (Fig. 4).

### The influence of life history on diversification

A popular perspective on diversification in this species complex is that paedomorphosis promotes reproductive isolation and, ultimately, speciation (10, 11). An important theme of our results is that taxa with different life history strategies commonly cluster together by geography in population genetic and phylogeographic analyses (Figs. 2, 3). Furthermore, genetic differentiation is not significantly greater for paedomorphic populations, relative to facultative populations, and there has been substantial gene flow among populations with different life history strategies (Fig. 4). Thus, the evolution of life history extremes does not appear to be the main driver of speciation in the tiger salamander complex.

It is also worth noting that facultative paedomorphosis is the inferred ancestral condition (8) and nearly all extant lineages in the *Ambystoma tigrinum* complex are capable of metamorphosis, either in the wild or in the lab. Only *A. dumerilii* is not known to survive metamorphosis, even under laboratory conditions (51). That only one described taxon in the complex is demonstrably fixed for paedomorphosis raises several questions: Are the rest of the primarily paedomorphic populations *en route* to fixation, rather than actually fixed? Have lineages fixed for paedomorphosis gone extinct? Do facultative populations fluctuate in paedomorph frequency over time, depending on environmental and/or demographic conditions (5)? Do transforming and non-transforming individuals interbreed in nature? While these remain open questions, below we provide some relevant context for exploration and future research.

Theory predicts that paedomorphosis in salamanders is most likely to become fixed in a population when ecological conditions are favorable in the larval, aquatic environment but unfavorable on land (e.g., when terrestrial habitats are arid or resource-limited) (12, 52, 53). However, it is not known how long fixed populations persist over evolutionary time; obligate paedomorphic lineages may have arisen and gone extinct multiple times over tiger salamander evolutionary history (54). One situation that would result in complete speciation is if paedomorphosis became fixed in the population and migration with surrounding facultatively paedomorphic populations ceased due to strong assortative mating and/or selection against hybrids (55–57). However, assortative mating within life history morphs has not been demonstrated in other salamanders (58–60) and different tiger salamander morphs can interbreed (61). In fact, the only known instance of positive assortative mating in *A. tigrinum* is associated with tail length (62, 63), which might favor metamorphosed males as paedomorphic individuals can have relatively short tails (42). The evidence for selection against intermorph offspring is also scarce. Captive breeding using artificial fertilization has produced *A. mexicanum* x *A. tigrinum*, *A. dumerilii* x *A. tigrinum*, *A. andersoni* x *A. tigrinum*, and *A. dumerilii* x *A. rivulare* hybrids (7, 31, 64–66). This suggests that postzygotic barriers are weak, although the extent to which divergent lineages would interbreed in the wild remains a key question in need of investigation.

If biological barriers to reproduction are limited or absent, there are several potential mechanisms by which dispersal could mediate gene flow between paedomorphic and metamorphic populations. First, metamorphosed individuals from facultatively paedomorphic populations could move into bodies of water containing paedomorphic populations. Our demographic analyses support this scenario, as they recovered immigration from facultative into paedomorphic populations (Fig. 4). However, demographic analyses also estimated emigration out of paedomorphic populations; thus, “obligate” paedomorphs might also occasionally metamorphose and disperse. Occasional metamorphosis has been documented in several putatively paedomorphic populations (9, 10, 42, 67–69), and may be more prevalent than currently recognized because rare transformed individuals in nature are virtually impossible to verify (70). Importantly, even a low frequency of transformed adults is likely sufficient to maintain evolutionary cohesion with surrounding populations (10, 29, 71). Alternatively, the dispersal of paedomorphs might be human-mediated; this could partially explain the elevated emigration rates we observed in the axolotl (Fig. 4). Axolotls are known to be sold in local markets as food, bait, or aquarium pets, and are sometimes moved from one area of central Mexico to another (72). Finally, gene flow may not necessarily be mediated across a terrestrial environment; genetic connectivity could also be maintained if populations have been forced into contact by water level fluctuations in the groundwater system of a region. The dynamic lacustrine history of the Cuenca Oriental (73), for instance, suggests that isolation of some paedomorphic populations has been punctuated by broader aquatic connections over short geological time scales, potentially facilitating gene flow. Collectively, these mechanisms may explain the maintenance of geographic genetic groups, now or in the evolutionarily recent past, containing a range of life histories.

The genetic cohesion we detected across wide swaths of the TMVB is particularly striking given other evidence for local adaptation in this region. This scenario is perhaps best exemplified in the Cuenca Oriental (Fig. 2E), where some lakes contain paedomorphic salamanders adapted to saline conditions, a very rare trait among amphibians (74). At one site, the saline Lake Alchichica, the population is currently considered a distinct species (*A. taylori*) (10). However, our results indicate that *A. taylori* is not genetically distinct from surrounding, putatively non-saline-adapted populations, a result consistent with a previous microsatellite-based study that found significant gene flow to and from *A. taylori* (29). While we cannot rule out the potential for localized selection and adaptation in the *A. taylori* genome, our results highlight that even populations adapted to aquatic conditions intolerable to most other *Ambystoma* have not reached a level of isolation identifying them as an independent evolutionary lineage (75).

### Comparisons to previous work

This study included extensive geographic sampling of the tiger salamander complex, which provided a broad spatial context to more fully understand patterns of genetic variation. However, it is important to consider the possibility that our data are simply not sensitive enough to detect genetic differentiation associated with lineage divergence on an extremely recent time scale. The common ancestor of *A. californiense* and the remainder of the species complex dates to approximately five million years (11, 34), and phylogenomic data indicate that speciation across the remaining tiger salamander lineages occurred within the last one million years (76). Such recent timing renders lineage boundaries difficult to detect (15). Our markers, developed from transcriptomic resources (32), may not have an overall substitution rate sufficient to detect such subtle genetic differentiation. We recommend that future research uses a larger genomic dataset or faster-evolving loci with particular focus on the sampling of paedomorphs in this system. To this point, perhaps the strongest evidence for the genetic divergence of paedomorphic populations comes from past microsatellite-based work indicating the genetic distinctiveness of *A. andersoni* and *A. mexicanum* from a select set of populations across central Mexico (27) and the genetic distinctiveness of *A. taylori* from neighboring populations in the Cuenca Oriental (29). Both of these studies recovered signatures of gene flow, but identified greater overall population structure compared to this study, which could be due to numerous factors, including a faster microsatellite mutation rate and more limited locality sampling.

## Conclusions

The extent to which populations within the tiger salamander complex exhibit phenotypic plasticity in life history traits is remarkable and is believed to have played a role in the rapid accumulation of lineages observed in the highlands of central Mexico (11, 57). While our results suggest there is less species-level diversity in central Mexico than previously recognized, there is clearly more diversity in this region compared to the US and Canada where there is (a) more geographic space and (b) less life history variation within and between lineages. While we cannot fully explain the greater diversity in central Mexico, our results suggest that major patterns of diversification are related to a complex history of geographic isolation and secondary contact, in which life history strategy has played a less important role, at least in the long (evolutionary) run. We agree with previous work (12) that the complex geological history of the TMVB, including montane uplift and fluctuating drainage connectivity since the Miocene, has been the cornerstone of the evolutionary history of this species complex, and that the influences of geographic isolation and paedomorphosis may work synergistically to lead to the establishment of isolated populations. Lingering questions notwithstanding, this large-scale genetic and geographic study establishes a framework for understanding the evolutionary history of the *Ambystoma tigrinum* species complex. The results presented here will facilitate comparative studies of the axolotl and its allies, provide direction for conservation prioritization and management, and strengthen the use of the tiger salamander species complex as a model system in biology.

## Materials & Methods

### Geographic sampling

We generated new data from 254 individuals sampled from across the range of the *A. tigrinum* complex (Fig. S1; Table S1). These individuals were combined with 93 individuals sampled in O’Neill et al. (32) to produce a data set comprising 347 individuals. We sampled a large number of localities (188) with 1-9 individuals sampled per locality (mean = 1.8). This sampling included 166 individuals from the US, 2 from Canada, and 178 from Mexico. *Ambystoma californiense* samples were primarily included as a close outgroup to the remainder of the species complex and were sampled from localities with limited to no impacts of introgression from invasive *A. mavortium* (47). Additional outgroup data were generated for two species outside of the *A. tigrinum* complex: *A. opacum* and *A. texanum*. Full details regarding the generation of these data can be found in the Supplementary Materials. All *A. tigrinum* complex species were assigned to a life history category based on review of the published literature and private field records from authors of this study. These life history categories are as follow: 1 = obligate metamorph (no paedomorphs have been documented in the field); 2 = strong bias towards metamorphosis with rare paedomorphs found in the field; 3 = both metamorphosis and paedomorphosis common in the field; 4 = strong bias towards paedomorphosis with rare metamorphs found in the field; 5 = obligate paedomorph (no metamorphs have been documented in the field). In the text, “paedomorph” describes taxa scored as 4 or 5, while “facultative paedomorph” describes taxa scored as 2 or 3.

### Data collection and sequencing

We generated DNA sequence data from a panel of 95 nuclear loci developed specifically for the tiger salamander species complex. A more complete description of marker development can be found in O’Neill et al. (32). Genomic DNA was extracted using a DNeasy Blood and Tissue kit (Qiagen). For a small number of DNA extractions, we increased DNA quantities using a Repli-g whole genome amplification kit (Qiagen). We used an initial round of PCR in 96-well plate format to amplify all loci from an individual, followed by a smaller second round of PCR to amplify loci that did not amplify in the initial PCR. See O’Neill et al. (32) for the details of PCR conditions and primer sequences.

PCR products from all loci were pooled for each individual in roughly equal concentrations based on the intensity of amplification as visualized on an agarose gel. Indexed Illumina sequencing libraries were generated for each individual using an Illumina Nextera XT DNA Library Preparation Kit (Illumina). Subsequent to library preparation, indexed libraries were quantified using a Qubit Fluorometer, pooled in equimolar concentrations, and checked on an Agilent Bioanalyzer to assure proper fragmentation.

Sequencing was performed in four rounds using an Illumina MiSeq. We performed an initial round using a total of seven individuals to test the compatibility of the Illumina Nextera XT library kit with our PCR amplicons. Three subsequent rounds of library preparation and sequencing were performed on sets of 96 individuals each, and some individuals were sequenced in multiple rounds due to initially low read counts. All sequencing was performed with paired-end 150 bp reads. Overall, we generated a total of 29,426,894 PE reads across all newly sequenced individuals, with an average of 309,757 PE reads, 1,148X coverage, and 7.6% missing loci per individual (Table S2).

### Bioinformatics and dataset generation

All sequence reads were processed using a newly developed bioinformatic pipeline written for this project (https://doi.org/10.5281/zenodo.3585970) that produces multiple sequence alignments for individual loci and genome-wide SNP matrices sampled from variable sites. This pipeline was developed using the Snakemake workflow management system (77), linking together multiple software tools to take sequence data from raw reads to phased sequence alignments for each locus. Demultiplexed paired-end Illumina fastq files were used as input, with separate forward and reverse read files for each individual. Sequence data were trimmed and filtered in Trimmomatic (78) using a sliding window of 4 base-pairs and a minimum average quality score of 15. Filtered sequence reads were then aligned to reference sequences from the O’Neill et al. (2013) dataset (32); specifically, we used the clean sequences of an *A. ordinarium* sample that had high coverage and low amounts of missing data. The resulting aligned contigs were processed using SAMtools (79) to filter and prepare data for FreeBayes (80), which was used to call variable sites. Variants were filtered with VCFTools (81) by removing indels and setting a quality threshold of phred score > 20 and a minimum read depth of 30. The program WhatsHap (82) was used to perform read-based phasing of the data for each locus x individual contig. Finally, phased haplotypes from each individual (two copies, regardless of homo- or heterozygosity) were combined into an alignment of all individuals using MAFFT with the default auto parameter (83). We generated fasta files of SNPs using the SNP-sites program (84) and created a SNP genotype matrix by sampling variable sites from a concatenated sequence alignment of all loci. We identified three loci (E12G1, E6A11, and E7G8) as potential paralogs based on high alignment error and high levels of heterozygosity for all individuals. These three loci were excluded from all analyses.

Across the remaining 92 loci, alignment lengths ranged from 123 to 630 bp (avg. = 270 bp) with a total concatenated alignment of 24,788 bp. For the full dataset, including *A. texanum* and *A. opacum* outgroups, single-locus alignments contained an average of 55 variable sites (min. = 15, max. = 121) and an average of 41 parsimony informative sites (min. = 11, max. = 102). Population genetic analyses restricted to the *A. tigrinum* complex including *A. californiense—*which were further filtered for non-biallelic SNPs and a minor allele count ≥ 3— contained a total of 2,360 SNPs.

### Population structure and lineage discovery

We developed hypotheses of population-level lineages across the range of the *A. tigrinum* complex ignoring the existing taxonomy, starting with the identification of major geographic patterns of differentiation, and then performing a recursive set of analyses on more geographically restricted sets of individuals. In our initial round of analyses we used two nonparametric methods: principal components analysis (PCA) and discriminant analysis of principal components (DAPC) (33). While both analyses provide a multivariate summary of genetic data, DAPC is also used to assess the fit of data to varying numbers of population clusters. These analyses were applied to our full genotypic dataset including *A. californiense* and all remaining individuals from the *A. tigrinum* complex. The PCA was calculated using the function ‘prcomp’ in the R package stats (85), while the DAPC was calculated using the package adegenet (86). The optimal number of principal components to retain for DAPC was identified using crossvalidation via the xvalDapc function with default parameter values. DAPC was performed without prior assignment of individuals to groups across a range of cluster levels (*K* = 1-20). We used two metrics to identify the best estimate of the primary splits in our data. First, we used the Bayesian Information Criterion (BIC) calculated in the DAPC analysis to assess the fit of the data to different levels of *K*. We note that the level of *K* with the absolute lowest BIC may not be a better explanation of the data than a *K* with a slightly higher BIC (87); therefore, we applied this measure for general guidance on a range of *K* that may describe the data well. We paired this assessment with visualizations of the first and second principal components (Fig. S2, S5) and DAPC ordination plots to identify the level of *K* at which similar clustering patterns could be observed with minimal change at successively higher levels of *K*. DAPC of the complete tiger salamander species complex identified a consistent pattern beginning at *K* = 5 for high differentiation of all *A. californiense* samples (Fig. S3). Further DAPC analysis with *A. californiense* removed identified a consistent pattern beginning at *K* = 5 for differentiation between clusters of populations from northern and central Mexico, and three clusters of U.S. populations (two from the Western U.S. and one from the Eastern U.S.; Fig. S4).

Using the clusters identified in the DAPC analysis of the total data set, we then used both DAPC and STRUCTURE v.2.3.4 (88) to analyze subsequent data sets comprising smaller numbers of individuals. Recursive rounds of DAPC analyses were stopped when BIC scores showed little improvement (ΔBIC < 2) at values of *K* > 1. STRUCTURE analyses used an admixture model and 500,000 generations following a burn-in of 100,000 generations. Analyses were performed for *K* = 1-10 with 16 replicate analyses for each *K*. To help identify an optimal value for *K*, we calculated *ΔK* using the Evanno method (89) via the CLUMPAK web tool (90). A limitation of the Evanno method is that it cannot estimate the likelihood of *K* = 1 (89), thus, we also visually inspected individual group assignments and concluded a value of *K* = 1 if the corresponding DAPC cluster showed little improvement (ΔBIC < 2) at values of *K* > 1, or when individuals were simply being split without any geographic or individual clustering. We also evaluated models where *K* was equal to the number of currently recognized species in that genetic cluster (e.g., *K* = 3 for CM1, which included the type localities for three described species: *A. altamirani*, *A. leorae*, and *A. rivulare*). We mapped individual species assignments onto these results to test the potential correspondence between naïve clustering and the existing taxonomy (Fig. S9).

### Phylogenetic reconstruction

Given the high levels of admixture among groups, we used SplitsTree v. 4.14.8 (35, 91) to generate four phylogenetic networks: one with all tiger salamander individuals, and one each for the U.S., central Mexico, and northern Mexico subgroups. Networks were constructed using uncorrected p-distances and the NeighborNet algorithm (92).

We also used three different analytical approaches to place hypothesized population lineages in a phylogenetic framework. For all analyses we used a reduced data set containing the concatenated data for 2-7 representative individuals from each population genetic cluster, which limited computation time and avoided violating the coalescent-model assumption of zero gene flow. We first inferred the phylogeny using Bayesian Inference in BEAST v.1.8.3 (37). Analyses were run for 5 million generations, sampling every 1,000 generations after the first 500,000 generations were removed as burn-in. Run convergence was assessed with Tracer v. 1.6.0 (93). Next, we inferred a maximum-likelihood phylogeny using RAxML v.8 (38). Node support was assessed using a rapid bootstrap analysis with 1,000 replicates, which was summarized as a 95% rule consensus tree using the program SumTrees in the DendroPy python library (94). For both BEAST and RAxML, PartitionFinder (95) was used to identify the number of preferred gene partitions and their substitution models, and analyses were performed on the CIPRES Science Gateway server (96). Finally, we inferred phylogenetic relationships using SVDquartets (36) implemented in PAUP* version 4.0a164 (97), sampling all possible quartets and assessing node support with 1,000 bootstrap replicates. For all phylogenetic analyses, trees were visualized using FigTree v. 1.4.2 (98).

### Tests of migration and population differentiation of paedomorphic taxa

We used the coalescent-based program Migrate-N v.3.2 (99) to explicitly test for population structure and gene flow in each obligately paedomorphic species of *Ambystoma* (*A. andersoni, A. dumerilii, A. taylori,* and *A. mexicanum).* For each model (Fig. S14), we treated paedomorphic species as distinct populations, and the facultatively paedomorphic individuals from the same genetic cluster (CM3 or CM4) as another population. The best-fitting model was determined via Bayes factors (BF), which were calculated using Bézier-corrected marginal likelihoods (Table S3); the highest-ranking models have a BF = 0. Each Migrate-N analyses ran for 50 million steps, recording every 10 steps, with a burn-in of five million and the default heating scheme. Suitable upper bounds for priors on population size (θ) and migration rate (*M*) were determined from an initial test run of 10 million steps for each analysis. Median values and 95% confidence intervals of θ and *M* are in Fig. 4 and Table S4.

Within CM3 and CM4, we also calculated pairwise Φ_ST_ (a metric of population differentiation) among all populations using the function ‘pairPhiST’ in the R package haplotypes (100). These values were plotted against pairwise geographic distances (Fig. 4), which were calculated from locality latitude/longitude data (Table S1) and converted to kilometers using the function ‘rdist.earth’ in the R package fields (101). A life history matrix was also generated by assigning a value of 0 for pairwise comparisons of populations with the same life history and a value of 1 for populations with different life histories (i.e., facultatively paedomorphic vs. obligately paedomorphic). Paedomorph-paedomorph comparisons were omitted to avoid confounding them with facultative-facultative comparisons. Finally, we used these matrices in a partial mantel test [function ‘mantel.partial’ package vegan (102)] to test for a correlation between Φ_ST_ and life history while correcting for geographic distance. Significance of this test was determined using 999 permutations and an alpha level of 0.05.

## Data Availability

Supplementary figures and tables are provided with the online version of this manuscript. Input files for all population genetic and phylogenetic analyses are available via figshare (https://figshare.com/projects/Life_history_strategy_does_not_reflect_genetic_differentiation_in_the_tiger_salamander_species_complex/74115). Sequence data are available on the NCBI Sequence Read Archive (BioProject accession PRJNA594660).

## Acknowledgements

This project was supported by grants from the National Science Foundation [DEB-0949532, DEB-1355000, and DEB-1406876 (DDIG awarded to J. D. Kratovil)] and by a University Research Fellowship awarded to K. M. Everson from the University of Kentucky. We thank Chris Beachy, Jim Bogart, Ken Carbale, Sheri Church, Jeff LeClere, Carol Hall, Paul Moler, Nancy Staub, and Ken Wray for genetic samples, and Jose Bocanegra, Ben Browning, Mackenzie Humphrey, Mary Virginia Gibbs, Ricky Grewelle, Alan Lemmon, Emily Moriarty Lemmon, Deborah Lu, Stephanie Mitchell, Alex Noble, Tolu Odukoya, Ben Tuttle, and Josh Williams for assistance with data collection and analyses.

## References

1. S. R. Palumbi, Marine speciation on a small planet. Trends Ecol. Evol. 7, 114–118 (1992).

2. S. Mopper, S. Y. Strauss, Genetic Structure and Local Adaptation in Natural Insect Populations: Effects of Ecology, Life History, and Behavior (Springer US, 1998).

3. S. J. Gould, Ontogeny and Phylogeny (Harvard University Press, 1977).

4. D. B. Wake, Paedomorphosis. J. Herpetol. 14, 80 (1980).

5. H. H. Whiteman, Evolution of facultative paedomorphosis in salamanders. Q. Rev. Biol. 69, 205–221 (1994).

6. R. B. Page, et al, Effect of thyroid hormone concentration on the transcriptional response underlying induced metamorphosis in the Mexican axolotl (Ambystoma). BMC Genomics 9, 78 (2008).

7. S. R. Voss, Genetic Basis of Paedomorphosis in the Axolotl, *Ambystoma mexicanum:* A Test of the Single-Gene Hypothesis. J. Hered. 86, 441–447 (1995).

8. S. R. Voss, H. B. Shaffer, Adaptive evolution via a major gene effect: paedomorphosis in the Mexican axolotl. Proc. Natl. Acad. Sci. USA 94, 14185–9 (1997).

9. E. H. Taylor, New salamanders from Mexico with a discussion of certain known forms. Univ. Kansas Sci. Bull. 26, 407–439 (1939).

10. R. Brandon, E. Maruska, W. Rumph, A new species of neotenic *Ambystoma* (Amphibia, Caudata) endemic to Laguna Alchichica, Puebla, Mexico. Bull. South. Calif. Acad. Sci. 80, 112–125 (1981).

11. H. B. Shaffer, M. L. McKnight, The polytypic species revisited: Genetic differentiation and molecular phylogenetics of the tiger salamander *Ambystoma tigrinum* (Amphibia: Caudata) complex. Evolution 50, 417–433 (1996).

12. H. B. Shaffer, Evolution in a paedomorphic lineage. I. An electrophoretic analysis of the Mexican ambystomatid salamanders. Evolution 38, 1194–1206 (1984).

13. H. B. Shaffer, Phylogenetics of model organisms: The laboratory axolotl, Ambystoma Mexicanum. Syst. Biol. 42, 508–522 (1993).

14. H. B. Shaffer, Evolution in a paedomorphic lineage. II. Allometry and form in the Mexican ambystomatid salamanders. Evolution 38, 1207–1218 (1984).

15. H. B. Shaffer, R. C. Thomson, Delimiting Species in Recent Radiations. Syst. Biol. 56, 896–906 (2007).

16. P. R. Grant, Speciation and the adaptive radiation of Darwin’s finches. Am. Sci. 69, 653–663 (1981).

17. O. Seehausen, Explosive speciation rates and unusual species richness in haplochromine cichlid fishes: Effects of sexual selection. Adv. Ecol. Res. 31, 237–274 (2000).

18. M. Fegan, P. Prior, “How complex is the *Ralstonia solanacearum* species complex” in Bacterial Wilt Disease and the Ralstonia Solanacearum Species Complex, C. Allen, P. Prior, A. C. Hayward, Eds. (American Phytopathological Society Press, 2005), pp. 449–461.

19. J. Mallet, Hybrid speciation. Nature 446, 279–283 (2007).

20. M. C. Fontaine, et al., Extensive introgression in a malaria vector species complex revealed by phylogenomics. Science 347, 1258524 (2015).

21. J. Geml, G. A. Laursen, K. O’Neill, H. C. Nusbaum, D. L. Taylor, Beringian origins and cryptic speciation events in the fly agaric (*Amanita muscaria*). Mol. Ecol. 15, 225–239 (2005).

22. R. Newman, Adaptive plasticity in amphibian metamorphosis. Bioscience 42, 671–678 (1992).

23. H. H. Whiteman, S. A. Wissinger, W. S. Brown, Growth and foraging consequences of facultative paedomorphosis in the tiger salamander, *Ambystoma tigrinum nebulosum*. Evol. Ecol. 10, 433–446 (1996).

24. P. J. Fernandez, J. P. Collins, Effect of environment and ontogeny on color pattern variation in Arizona tiger salamanders (*Ambystoma tigrinum nebulosum* Hallowell). Copeia 1988, 928 (1988).

25. S. P. D. Riley, H. B. Shaffer, S. R. Voss, B. M. Fitzpatrick, Hybridization between a rare, native tiger salamander (*Ambystoma californiense*) and its introduced congener. Ecol. Appl. 13, 1263–1275 (2003).

26. D. W. Weisrock, H. B. Shaffer, B. L. Storz, S. R. Storz, S. R. Voss, Multiple nuclear gene sequences identify phylogenetic species boundaries in the rapidly radiating clade of Mexican ambystomatid salamanders. Mol. Ecol. 15, 2489–2503 (2006).

27. G. Parra-Olea, et al., Conservation genetics of threatened Mexican axolotls (*Ambystoma*). Anim. Conserv. 15, 61–72 (2012).

28. E. Routman, Population structure and genetic diversity of metamorphic and paedomorphic populations of the tiger salamander, *Ambystoma tigrinum*. J. Evol. Biol. 6, 329–357 (1993).

29. R. Percino-Daniel, E. Recuero, E. Vazquez-Dominguez, K. R. Zamudio, G. Parra-Olea, All grown-up and nowhere to go: Paedomorphosis and local adaptation in *Ambystoma* salamanders in the Cuenca Oriental of Mexico. Biol. J. Linn. Soc. 118, 582–597 (2016).

30. R. A. Brandon, Hybridization between the Mexican salamanders *Ambystoma dumerilii* and *Ambystoma mexicanum* under laboratory conditions. Herpetologica 28, 199–207 (1972).

31. R. A. Brandon, Interspecific hybridization among Mexican and United States salamanders of the genus *Ambystoma* under laboratory conditions. Herpetologica 33, 133–152 (1977).

32. E. M. O’Neill, et al., Parallel tagged amplicon sequencing reveals major lineages and phylogenetic structure in the North American tiger salamander (*Ambystoma tigrinum*) species complex. Mol. Ecol. 22, 111–129 (2013).

33. T. Jombart, S. Devillard, F. Balloux, Discriminant analysis of principal components: A new method for the analysis of genetically structured populations. BMC Genet. 11, 94 (2010).

34. H. B. Shaffer, G. B. Pauly, J. C. Oliver, P. C. Trenham, The molecular phylogenetics of endangerment: Cryptic variation and historical phylogeography of the California tiger salamander, *Ambystoma californiense*. Mol. Ecol. 13, 3033–3049 (2004).

35. D. H. Huson, SplitsTree: Analyzing and visualizing evolutionary data. Bioinformatics 14, 68–73 (1998).

36. J. Chifman, L. Kubatko, Quartet inference from SNP data under the coalescent model. Bioinformatics 30, 3317–3324 (2014).

37. A. J. Drummond, A. Rambaut, BEAST: Bayesian evolutionary analysis by sampling trees. BMC Evol. Biol. 7, 214 (2007).

38. A. Stamatakis, RAxML version 8: A tool for phylogenetic analysis and post-analysis of large phylogenies. Bioinformatics 30, 1312–1313 (2014).

39. A. A. Dugès, Description d’un axolotl des montagnes de las cruces (Amblystoma [sic] altamirani, A. Dugès) (Imprimerie du Ministère de “Fomento,” 1895).

40. E. H. Taylor, Herpetological novelties from Mexico. Univ. Kansas Sci. Bull. 29, 343–358 (1943).

41. E. H. Taylor, A new ambystomid salamander from the Plateau Region of Mexico. Univ. Kansas Sci. Bull. 30, 57–61 (1944).

42. S. Krebs, R. Brandon, A new species of salamander (family Ambystomatidae) from Michoacan, Mexico. Herpetologica 40, 238–245 (1984).

43. A. A. Dugès, Una nueva especie de ajolote de la Laguna de Patzcuaro. La Nat. 1, 241–244 (1870).

44. J. Dixon, A new species of salamander of the genus Ambystoma from Jalisco, Mexico. Copeia 1, 99–101 (1963).

45. G. Shaw, F. Nodder, The Naturalist’s Miscellany; or Coloured Figures of Natural Objects Drawn and Described Immediately from Nature, 9th Ed. (Nodder & Co., 1798).

46. A. A. Dugès, Erpetología del valle de México. La Nat. 2, 97–146 (1888).

47. J. R. Johnson, R. C. Thomson, S. J. Micheletti, H. B. Shaffer, The origin of tiger salamander (*Ambystoma tigrinum*) populations in California, Oregon, and Nevada: Introductions or relicts? Conserv. Genet. 12, 355–370 (2011).

48. E. H. Taylor, Two new ambystomid salamanders from Chihuahua. Copeia 3, 143–146 (1941).

49. R. Webb, Observations on tiger salamanders (*Ambystoma tigrinum* complex, family Ambystomatidae) in Mexico with description of a new species. Bull. Maryl. Herpetol. Soc. 40, 122–143 (2004).

50. H. B. Shaffer, Biosystematics of *Ambystoma rosaceum* and *A. tigrinum* in Northwestern Mexico. Copeia 1983, 67 (1983).

51. R. A. Brandon, Spontaneous and induced metamorphosis of *Ambystoma dumerilii* (Dugès), a paedogenetic Mexican salamander, under laboratory conditions. Herpetologica 32, 429–438 (1976).

52. H. M. Wilbur, J. P. Collins, Ecological aspects of amphibian metamorphosis: Nonnormal distributions of competitive ability reflect selection for facultative metamorphosis. Science 182, 1305–1314 (1973).

53. R. M. Bonett, M. A. Steffen, S. M. Lambert, J. J. Wiens, P. T. Chippindale, Evolution of paedomorphosis in plethodontid salamanders: Ecological correlates and re-evolution of metamorphosis. Evolution 68, 466–482 (2014).

54. C. R. Marshall, E. C. Raff, R. A. Raff, Dollo’s law and the death and resurrection of genes. Proc. Natl. Acad. Sci. USA 91, 12283–12287 (1994).

55. M. J. West-Eberhard, Phenotypic plasticity and the origins of diversity. Annu. Rev. Ecol. Syst. 20, 249–278 (1989).

56. M. J. West-Eberhard, Alternative adaptations, speciation, and phylogeny (A Review). Proc. Natl. Acad. Sci. USA 83, 1388–92 (1986).

57. B. M. Fitzpatrick, Underappreciated consequences of phenotypic plasticity for ecological speciation. Int. J. Ecol. 2012, 1–12 (2012).

58. M. Denoël, P. Poncin, J.-C. Ruwet, Sexual compatibility between two heterochronic morphs in the Alpine newt, *Triturus alpestris*. Anim. Behav. 62, 559–566 (2001).

59. J. D. Krenz, P. A. Verrell, Integrity in the midst of sympatry: Does sexual incompatibility facilitate the coexistence of metamorphic and paedomorphic mole salamanders (*Ambystoma talpoideum)?* J. Zool. 258, S0952836902001589 (2002).

60. H. H. Whiteman, J. D. Krenz, R. D. Semlitsch, Intermorph breeding and the potential for reproductive isolation in polymorphic mole salamanders (*Ambystoma talpoideum*). Behav. Ecol. Sociobiol. 60, 52–61 (2006).

61. H. Whiteman, J. Gutrich, R. Moorman, Courtship behavior in a polymorphic population of the tiger salamander, *Ambystoma tigrinum nebulosum*. J. Herpetol. 33, 348–351 (1999).

62. R. D. Howard, R. S. Moorman, H. H. Whiteman, Differential effects of mate competition and mate choice on eastern tiger salamanders. Anim. Behav. 53, 1345–1356 (1997).

63. D. H. Bos, R. N. Williams, D. Gopurenko, Z. Bulut, J. A. DeWoody, Condition-dependent mate choice and a reproductive disadvantage for MHC-divergent male tiger salamanders. Mol. Ecol. 18, 3307–3315 (2009).

64. C. E. Nelson, R. R. Humphrey, Artificial interspecific hybridization among *Ambystoma*. Herpetologica 28, 27–32 (1972).

65. S. R. Voss, H. B. Shaffer, Evolutionary genetics of metamorphic failure using wild-caught vs. laboratory axolotls (*Ambystoma mexicanum*). Mol. Ecol. 9, 1401–1407 (2000).

66. S. R. Voss, K. L. Prudic, J. C. Oliver, H. B. Shaffer, Candidate gene analysis of metamorphic timing in ambystomatid salamanders. Mol. Ecol. 12, 1217–1223 (2003).

67. J. B. Armstrong, G. M. Malacinski, Developmental Biology of the Axolotl (Oxford University Press, 1989).

68. J. M. Velasco, Anotaciones y observaciones al trabajo del Sr. D. A. Weismann sobre la transformación del ajolote Mexicano en Amblistoma. La Nat. 5, 58–84 (1880).

69. H. M. Smith, The Mexican axolotl: Some misconceptions and problems. Bioscience 19, 593–615 (1969).

70. L. L. Bailey, T. R. Simons, K. H. Pollock, Estimating site occupancy and species detection probability parameters for terrestrial salamanders. Ecol. Appl. 14, 692–702 (2004).

71. C. L. Morjan, L. H. Rieseberg, How species evolve collectively: Implications of gene flow and selection for the spread of advantageous alleles. Mol. Ecol. 13, 1341–1356 (2004).

72. J. McKay, “An evaluation of captive breeding and sustainable use of the Mexican axolotl (*Ambystoma mexicanum*),” University of Kent, Canterbury, UK. (2003) (January 20, 2020).

73. J. L. Tamayo, R. C. West, “The hydrography of middle America” in Handbook of Middle American Indians, R. Wauchope, R. C. West, Eds. (University of Texas Press, 1964), pp. 84–121.

74. G. R. Hopkins, E. D. Brodie, Occurrence of amphibians in saline habitats: A review and evolutionary perspective. Herpetol. Monogr. 29, 1–27 (2015).

75. K. De Queiroz, Species concepts and species delimitation. Syst. Biol. 56, 879–886 (2007).

76. P. M. Hime, et al., Phylogenomics reveals ancient gene tree discordance in the amphibian tree of life. Syst. Biol. (2020) https://doi.org/10.1093/sysbio/syaa034 (May 12, 2020).

77. J. Koster, S. Rahmann, Snakemake: A scalable bioinformatics workflow engine. Bioinformatics 28, 2520–2522 (2012).

78. A. M. Bolger, M. Lohse, B. Usadel, Trimmomatic: A flexible trimmer for Illumina sequence data. Bioinformatics 30, 2114–2120 (2014).

79. H. Li, A statistical framework for SNP calling, mutation discovery, association mapping and population genetical parameter estimation from sequencing data. Bioinformatics 27, 2987–2993 (2011).

80. E. Garrison, G. Marth, Haplotype-based variant detection from short-read sequencing. arXiv 1207.3907 (2012).

81. P. Danecek, et al., The variant call format and VCFtools. Bioinformatics 27, 2156–2158 (2011).

82. M. Martin, et al., WhatsHap: Fast and accurate read-based phasing. bioRxiv 10.1101/08 (2016).

83. K. Katoh, D. M. Standley, MAFFT multiple sequence alignment software version 7: Improvements in performance and usability. Mol. Biol. Evol. 30, 772–780 (2013).

84. A. J. Page, et al., SNP-sites: Rapid efficient extraction of SNPs from multi-FASTA alignments. Microb. genomics 2, e000056 (2016).

85. R Core Team, R: a language and environment for statistical computing (2012).

86. T. Jombart, adegenet: A R package for the multivariate analysis of genetic markers. Bioinformatics 24, 1403–1405 (2008).

87. T. Jombart, C. Collins, A tutorial for discriminant analysis of principal components (DAPC) using adegenet 2.0. 0. adegenet.r-forge.r-project.org (2015) (May 30, 2019).

88. J. K. J. Pritchard, M. Stephens, P. Donnelly, Inference of population structure using multilocus genotype data. Genetics 155, 945–959 (2000).

89. G. Evanno, S. Regnaut, J. Goudet, Detecting the number of clusters of individuals using the software structure: A simulation study. Mol. Ecol. 14, 2611–2620 (2005).

90. N. M. Kopelman, J. Mayzel, M. Jakobsson, N. A. Rosenberg, I. Mayrose, Clumpak: A program for identifying clustering modes and packaging population structure inferences across K. Mol. Ecol. Resour. 15, 1179–1191 (2015).

91. D. H. Huson, D. Bryant, Application of phylogenetic networks in evolutionary studies. Mol. Biol. Evol. 23, 254–267 (2006).

92. D. Bryant, V. Moulton, Neighbor-Net: An agglomerative method for the construction of phylogenetic networks. Mol. Biol. Evol. 21, 255–265 (2003).

93. A. Rambaut, A. Drummond, D. Xie, G. Baele, M. Suchard, Tracer v1.7 (2018).

94. J. Sukumaran, M. Holder, DendroPy: A Python library for phylogenetic computing. Bioinformatics 26, 1569–1571 (2010).

95. R. Lanfear, B. Calcott, S. Y. W. Ho, S. Guindon, PartitionFinder: Combined selection of partitioning schemes and substitution models for phylogenetic analyses. Mol Biol Evol 29, 1695–1701 (2012).

96. M. A. Miller, W. Pfeiffer, T. Schwartz, Creating the CIPRES Science Gateway for inference of large phylogenetic trees in 2010 Gateway Computing Environments Workshop (GCE), (IEEE, 2010), pp. 1–8.

97. D. Swofford, PAUP 4.0 b10: Phylogenetic analysis using parsimony (2002) (December 21, 2013).

98. A. Rambaut, FigTree, a graphical viewer of phylogenetic trees (2007) (May 8, 2014).

99. P. Beerli, M. Palczewski, Unified framework to evaluate panmixia and migration direction among multiple sampling locations. Genetics 185, 313–326 (2010).

100. C. Aktas, Haplotypes: Haplotype inference and statistical analysis of genetic variation (2015).

101. D. Nychka, R. Furrer, J. Paige, S. Sain, fields: Tools for spatial data (2017).

102. P. Dixon, VEGAN, a package of R functions for community ecology. J. Veg. Sci. 14, 927–930 (2003).

