## Supplementary Materials for "Geography is more important than life history in the recent diversification of the tiger salamander complex"

PNAS
www.pnas.org

### Supplementary Information for

#### Geography is more important than life history in the recent diversification of the tiger salamander complex

Kathryn M. Everson*, Levi N. Gray*, Angela G. Jones, Nicolette M. Lawrence, Mary E. Foley, Kelly L. Sovacool, Justin D. Kratovil, Scott Hotaling, Paul M. Hime, Andrew Storfer, Gabriela Parra-Olea, Ruth Percino-Daniel, X. Aguilar-Miguel, Eric. M. O'Neill, Luis Zambrano, H. Bradley Shaffer, David W. Weisrock

*Co-first authors
Corresponding author: David W. Weisrock


**This PDF file includes:**

Supplementary text
Figs. S1 to S10
Tables S1 to S3
References for SI citations

#### Supplementary Text

*Taxonomic recommendations for the* A. tigrinum *species complex.*

Our goal in the following is to make taxonomic recommendations consistent with our findings for the *Ambystoma tigrinum* complex. We provide an updated framework within which future studies can operate until lineage boundaries are further refined with additional ecological, morphological, and genomic data, and with reference to type specimens and series. We also take a conservative approach; in cases where obligate (or near obligate) paedomorphic populations are not genetically distinguishable with our data from surrounding facultatively paedomorphic populations, we recommend the use of subspecies. We find it useful to recognize biologically meaningful variation below the species level for a number of practical reasons (1). Locally adapted populations, for instance, can play important roles in the evolution of a lineage (2–4). There are likely evolutionary consequences for variation in frequency of paedomorphosis among populations, which we hope will be investigated further in the tiger salamander complex (see, e.g. 5). Characterizing fine-scale patterns of morphological diversity among populations in this complex would be an important contribution, particularly with respect to the genetic groups identified in this study.

The CM1 group included individuals that could be assigned to three recognized species: *Ambystoma* [=*Amblystoma*] *altamirani* Duges 1895 (6)*, A. leorae* (Taylor 1943) (7)*,* and *A. rivulare* [=*A. rivularis* (Taylor 1940)] (8). These three taxa (*A. altamirani, A. leorae,* and *A. rivulare*) are facultatively paedomorphic species that primarily occupy clear, fast-flowing streams. These taxa were placed in a distinct genus in the past, *Rhyacosiredon* Dunn 1928 (9). All have the same general coloration and are difficult to distinguish morphologically; Taylor's (7, 8) diagnoses focused primarily on subtle differences in body length and head shape. Within CM1, our STRUCTURE analysis identified $K = 2$, with *A. rivulare* in a western geographic group and the other two taxa in an eastern group; however, we note that these groups are genetically admixed (Fig. 2). If we consider admixture as evidence that these groups are not reproductively isolated, a single species name could apply to all of these populations: following rules of taxonomic priority, the names *A. leorae* and *A. rivulare* should be synonymized with the oldest name, *A. altamirani*. However, given that fine-scale genetic differentiation is present in this group (S7, S8a), we recommend additional investigation using multiple lines of evidence to clarify species limits in CM1. Due to potential geographic isolation and limited admixture with the other populations when assessing CM1 at $K = 3$ (S8a), we recommend use of a subspecific epithet for populations in the Rio Frio region (*A. altamirani leorae*).

CM2 is also composed of geographically proximate populations representing three named forms: *A. bombypellum* [=*A. bombypella* Taylor 1939] (10), *A. granulosum* Taylor 1944 (11), and *A. lermaense* [=*Siredon lermaensis* (Taylor 1939)] (10). Both DAPC and STRUCTURE results yielded strong support for a single cluster, and all individuals used in our phylogenetic analyses were recovered as a single monophyletic group [RaxML bootstrap support (BS) = 100; BEAST Bayesian posterior probability (BPP) = 0.95; SVDquartets BS = 100]. We interpret these results as strong evidence that a single species in this group should be recognized. We suggest that *A. granulosum* and *A. bombypellum* should be synonymized and assigned to the single species name *A. lermaense.*

The CM3 group contains individuals from multiple recognized species with a variety of life history strategies: three facultatively paedomorphic taxa [*A.*

*amblycephalum* [=*A. amblycephala* Taylor 1939] (10), *A. flavipiperatum* Dixon 1963 (12), and *A. ordinarium* [=*A. ordinaria* Taylor 1939] (10) and two taxa that are considered obligate paedomorphs [*A. andersoni* Krebs & Brandon 1984 (13) and *A. dumerilii* (Duges 1870) (14); but see (13, 15) for descriptions of occasional metamorphosed individuals under laboratory conditions]. A more exclusive STRUCTURE analysis of this group revealed that *A. ordinarium* does form a cluster that is genetically distinct from the remaining CM3 populations; this result was confirmed by phylogenetic analyses and corroborated by other studies (16, 17). Furthermore, *A. ordinarium* is phenotypically distinct from the remaining members of the CM3 clade (18). We did not recover any evidence for a second, cryptic species in the western portion of *A. ordinarium*'s range, which was hypothesized in a previous study (17). We recommend continuing to recognize *A. ordinarium* a single, distinct species, although we recommend future fine-scale analyses with a larger marker set to help understand its evolutionary history in the group.

The remaining species belonging to CM3 (*A. amblycephalum*, *A. flavipiperatum*, *A. andersoni*, and *A. dumerilii*) are highly admixed, which might be interpreted as "good" news for these highly threatened taxa, as their overall population sizes and geographic ranges might actually be larger than previously recognized (Fig S8c). However, we did find evidence for reproductive isolation among the obligate paedomorphic taxa, *A. andersoni* and *A. dumerilii*. We emphasize that *A. dumerilii*, despite showing substantial admixture with other populations in CM3, is still an ecologically unique, locally adapted lineage that warrants additional research and continued protection. As the only fixed paedomorphic lineage, further investigation into the timing and divergence of this population should attempt to determine the geographic context of speciation. We also recommend continued usage of *A. andersoni* Krebs & Brandon 1984 (13) for the same reasons. However, we note that none of our comparative samples were in close geographic proximity to Lake Zacapú (the nearest sample was collected from a locality 43 kilometers away) so additional work is needed to verify whether this species is truly isolated from nearby transforming populations. Finally, we recommend that *A. flavipiperatum* and *A. amblycephalum* be recognized by a single species name, with *A. amblycephalum* having priority.

Our results indicate that the Mexican axolotl, *A. mexicanum* (Shaw & Nodder 1798) (19), should continue to be recognized as a distinct species; however, the remaining members of CM4 represent a more challenging taxonomic scenario. Taylor (20) originally described the species *A. subsalsum* in CM4's range – the Lake Alchichica area – using a field-caught, metamorphosed individual as the type specimen and paedomorphic/aquatic individuals to fill out the type series. Later, Brandon et al. (21) made two taxonomic changes: (1) all metamorphic individuals assigned to the name *A. subsalsum* were synonymized with *A. tigrinum velasci* Green 1825 (22) (= *A. velasci*), and (2) the aquatic individuals in Lake Alchichica were described as a new species, *A. taylori* Brandon et al. 1981 (21). Our results showed a single genetic group in the Cuenca Oriental region made up of both *A. taylori* and *A. velasci*. Our results therefore appear to support Taylor's original decision to describe *A. subsalsum* from the Alchichica area using both metamorphic and paedomorphic/aquatic individuals in the type series. Despite this finding, we cannot rule out the possibility that this lineage also occurred (and still occurs) in the vicinity of Mexico City, where *A. velasci* is described from and has taxonomic priority. Other evidence exists for two species of *Ambystoma* occurring in the Xochimilco area of Mexico City (23–26). We therefore recommend use of the name *A. velasci* Green

1825 (22) for this other species and suggest follow-up studies be done to locate transformed individuals in the vicinity of Mexico City and to verify overall distribution. Attempts could also be made to extract DNA from the type series of *A. velasci* to verify the genetic distinctiveness of those individuals from the Mexico City area. If specimens from the type series of *A. velasci* turn out to be genetically indistinguishable from *A. mexicanum*, which is known to transform occasionally, *A. velasci* should be synonymized with *A. mexicanum* and the name *A. subsalsum* would have priority for the other CM4 lineage. We continue to recognize that the aquatic Alchichica population is unique in being adapted to levels of salinity that most amphibians would not be able to tolerate (10, 27); thus, we recommend a subspecific epithet (*A. velasci taylori*) for individuals in this lake.

We identified two individuals that were collected well outside the range of their assigned group. DWW-3104 and DWW-1686 were assigned to the CM2/*A. bombypellum* group, yet were collected within the ranges of CM4 and CM3, respectively. Pending additional work, we assume that these represent errors in specimen labeling or data transcription during the collection or lab-work stages.

Within the U.S. and northern Mexico, we recommend relatively few changes to the current taxonomy. Across the U.S. and Canada, as many as six taxa (species and subspecies) are currently recognized. If one were to assign taxonomic names corresponding to the eastern, central, and Rocky Mountain clades we recovered, these would be *Ambystoma tigrinum* Green 1825 (22), *A. mavortium mavortium* Baird 1850 (28), and *A. m. nebulosum* Hallowell 1853 (29), respectively. We agree with other recent authors that *A. mavortium* should continue to be considered a full species (it was resurrected from synonymy with *A. tigrinum*; 21). *Ambystoma tigrinum* and *A. mavortium* have several phenotypic and life history differences (30, 31), including a near-absence of paedomorphosis in *A. tigrinum*. We also found evidence that *A. m. nebulosum* should continue to be recognized as a valid subspecies, as it forms a monophyletic group and it occupies a somewhat distinct high-elevation habitat, despite showing substantial admixture with the remaining individuals from the *A. mavortium* cluster. However, we found limited evidence for other described subspecies. Future work using genetic data specifically suited for fine-scale population genetics will be needed to determine the validity and geographic ranges of *A. m. stebbinsi, A. m. diaboli,* and *A. m. melanostictum*.

The two groups recovered in our northern Mexico analyses most likely correspond to *A. rosaceum* Taylor 1941 (32) and *A. silvense* Webb 2004 (33), but as we were not able to collect any individuals directly from the type locality of *A. silvense*, we cannot be certain in that assignment. Salamanders are also thought to occur throughout northern Mexico in the areas directly east of our sampling range (e.g., see range maps for *A. tigrinum* and *A. velasci* at iucnredlist.org); however, without having genetic data from individuals in that region, we cannot pinpoint where the geographic boundaries of the north Mexican taxa lie, nor where the most admixture is occurring. Notably, Webb (33) used the name *A. subsalsum* for several specimens from eastern Durango and other parts of the Mexican Plateau (albeit "provisionally," p. 126); however, we do not believe that the name *A. subsalsum* should be associated with any specimens collected in that region. Some of those populations Webb assigned as *A. subsalsum* we assign as members of CM3 (*A. amblycephalum*). See the paragraph related to CM4 above for our discussion of the name *A. subsalsum*.

*Species assignments for STRUCTURE comparisons in supplement*

To visually assess how well current taxonomy matches our population genetics results, we assigned species names to samples *a posteriori* using a combination of our results, published literature, and geographic information. In some cases, these assignments would not match up with what biologists would assign to individuals in the field *a priori*. For instance, our study demonstrates that samples historically assigned as *A. velasci* in Nuevo Leon, Guanajuato, and San Luis Potosi fall into CM3 rather than CM4 where they would be expected. In that case, we assigned them as *A. flavipiperatum* because all of our analyses place those samples in CM3 and the localities are closer to the *A. flavipiperatum* type locality than to *A. amblycephalum*. Below we describe details of the more complicated assignments within CM groups.

Within CM1, records for *Ambystoma altamirani* and *A. rivulare* overlap in geographic space. We resolved this by assigning samples from west of Toluca, Mexico as *A. rivulare*.

Records for all three species represented in CM2 are mostly overlapping. We assigned samples from near the type locality of *Ambystoma lermaense* ("Lake Lerma, east of Toluca, México") as *A. lermaense*, and records from above 19.6 degrees N latitude as *A. bombypellum*. All other records in this group were assigned as *A. granulosum*, which was expected to have a larger distribution than the others.

Within CM3, only *Ambystoma amblycephalum* and *A. flavipiperatum* posed a challenge. Since *A. flavipiperatum* was believed to have a very small distribution near the type locality, we only assigned records near Santa Cruz, Jalisco as representatives of that species. All other samples were assigned to *A. amblycephalum*, which was reported to have a broader distribution than *A. flavipiperatum*.

For CM4, both *Ambystoma mexicanum* and *A. taylori* are paedomorphs with small distributions in Mexico City and Lake Alchichica, respectively. All other records were assigned to *A. velasci*.

*Generation of sequence data for phylogenetic outgroups.*

We retrieved sequence data from two outgroup taxa (*A. talpoidium* and *A. opacum*) using whole genome data published by Hime et al. (34). The *de novo* genome assemblies were set as custom BLAST (NCBI) databases in Geneious v.6.1.8 (35), and orthologous loci were retrieved by searching the custom databases for the 92 probe sequences used in this study. All search hits longer than 100 nucleotides with > 80% similarity were pulled from the assembly data, aligned as described in the main text, and trimmed to match the sequence length of the remaining data matrix.

Two additional outgroup individuals (DWW3233 and DWW2561) were sequenced with the primary dataset. At the time of collection these individuals were presumed to belong to the tiger salamander species complex, but preliminary results revealed them to be genetically distinct; thus, we sequenced a mitochondrial barcoding gene [NADH dehydrogenase subunit 2 (ND2)] to confirm species identity. Amplifications were performed in 30 μL reactions containing amplification buffer, Taq polymerase, dNTPs, purified water, template DNA, forward primer L4437 (5'-AAGCTTTCGGGCCCATACC-3'), and reverse primer H5692 (5'-GCGTTTAGCTGTTAACTAAA-3'). PCR thermal cycling conditions were initial denaturation at 94 °C for 3 min, followed by 30 cycles of 94°C for 30 s, 50°C for 45 s, and 72°C for 90 s, and a final extension of 72 °C for 5 min. Aliquots of the PCR products were electrophoresed and visualized on 1% agarose gels, and were purified and sequenced by Eurofins Genomics (Louisville, Kentucky, USA). The resulting ND2 sequences were compared to sequences in the NCBI BLAST database (http://blast.ncbi.nlm.nih.gov/Blast.cgi). The taxonomic identity of DWW3233 matched *A. texanum* with ≥ 99% sequence similarity, while DWW2561 matched *A. opacum* with ≥ 99% similarity.

#### Supplementary References


1. K. Winker, Chapter 1: Subspecies represent geographically partitioned variation, a gold mine of evolutionary biology, and a challenge for conservation. *Ornithol. Monogr.* **67**, 6–23 (2010).
2. E. Sanford, M. W. Kelly, Local Adaptation in Marine Invertebrates. *Ann. Rev. Mar. Sci.* **3**, 509–535 (2011).
3. R. I. Colautti, J. A. Lau, Contemporary evolution during invasion: evidence for differentiation, natural selection, and local adaptation. *Mol. Ecol.* **24**, 1999–2017 (2015).
4. V. Ravigné, U. Dieckmann, I. Olivieri, Live where you thrive: joint evolution of habitat choice and local adaptation facilitates specialization and promotes diversity. *Am. Nat.* **174**, E141-69 (2009).
5. H. H. Whiteman, Evolution of facultative paedomorphosis in salamanders. *Q. Rev. Biol.* **69**, 205–221 (1994).
6. A. A. Dugès, *Description d'un axolotl des montagnes de las cruces (*Amblystoma *[sic]* altamirani*, A. Dugès)* (Imprimerie du Ministère de "Fomento," 1895).
7. E. H. Taylor, Herpetological novelties from Mexico. *Univ. Kansas Sci. Bull.* **29**, 343–358 (1943).
8. E. H. Taylor, A new *Rhyacosiredon* (Caudata) from Western Mexico. *Herpetologica* **1**, 171–176 (1940).
9. E. Dunn, A new genus of salamanders from Mexico. *Proc. New Engl. Zool. Club* **10**, 85–86 (1928).
10. E. H. Taylor, New salamanders from Mexico with a discussion of certain known forms. *Univ. Kansas Sci. Bull.* **26**, 407–439 (1939).
11. E. H. Taylor, A new ambystomid salamander from the Plateau Region of Mexico. *Univ. Kansas Sci. Bull.* **30**, 57–61 (1944).
12. J. Dixon, A new species of salamander of the genus Ambystoma from Jalisco, Mexico. *Copeia* **1**, 99–101 (1963).
13. S. Krebs, R. Brandon, A new species of salamander (family Ambystomatidae) from Michoacan, Mexico. *Herpetologica* **40**, 238–245 (1984).
14. A. A. Dugès, Una nueva especie de ajolote de la Laguna de Patzcuaro. *La Nat.* **1**, 241–244 (1870).
15. R. A. Brandon, Spontaneous and induced metamorphosis of *Ambystoma dumerilii* (Dugès), a paedogenetic Mexican salamander, under laboratory conditions. *Herpetologica* **32**, 429–438 (1976).
16. D. W. Weisrock, H. B. Shaffer, B. L. Storz, S. R. Storz, S. R. Voss, Multiple nuclear gene sequences identify phylogenetic species boundaries in the rapidly radiating clade of Mexican ambystomatid salamanders. *Mol. Ecol.* **15**, 2489–2503 (2006).
17. P. M. Hime, *et al.*, The influence of locus number and information content on species delimitation: An empirical test case in an endangered Mexican salamander. *Mol. Ecol.* **25**, 5959–5974 (2016).
18. J. D. Anderson, R. D. Worthington, The life history of the Mexican salamander *Ambystoma ordinarium* Taylor. *Herpetologica* **27**, 165–176 (1971).

19. G. Shaw, F. Nodder, *The Naturalist's Miscellany; or Coloured Figures of Natural Objects Drawn and Described Immediately from Nature*, 9th Ed. (Nodder & Co., 1798).
20. E. H. Taylor, A New Ambystomid Salamander Adapted to Brackish Water. *Copeia* **1943**, 151 (1943).
21. R. Brandon, E. Maruska, W. Rumph, A new species of neotenic *Ambystoma* (Amphibia, Caudata) endemic to Laguna Alchichica, Puebla, Mexico. *Bull. South. Calif. Acad. Sci.* **80**, 112–125 (1981).
22. J. Green, Description of a new species of salamander. *J. Acad. Nat. Sci.* **5**, 116–118 (1825).
23. H. M. Smith, R. B. Smith, "Analysis of the literature on the Mexican axolotl" in *Synopsis of the Herpetofauna of Mexico, v. 1*, (E. Lundberg, 1971), pp. 1–245.
24. R. A. Brandon, "Natural history of the axolotl and its relationship to other ambystomatid salamanders" in *Developmental Biology of the Axolotl*, J. B. Armstrong, G. M. Malacinski, Eds. (Oxford University Press, 1989), pp. 13–21.
25. H. M. Smith, The Mexican axolotl: some misconceptions and problems. *Bioscience* **19**, 593–615 (1969).
26. J. M. Velasco, Anotaciones y observaciones al trabajo del Señor Augusto Weismann sobre la transformacion del ajolote Mexicano en *Amblistoma*. *La Nat.* **5**, 58–84 (1880).
27. V. Shoemaker, K. A. Nagy, Osmoregulation in amphibians and reptiles. *Annu. Rev. Physiol.* **39**, 449–471 (1977).
28. S. Baird, Revision of the North American Tailed-Batrachia with descriptions of new genera and species. *J. Acad. Nat. Sci. Philadelphia* **2**, 281–294 (1850).
29. E. Hallowell, On some new reptiles from California. *Proc. Natl. Acad. Sci. Philadelphia* **6**, 236–238 (1853).
30. D. Irschick, H. Shaffer, The polytypic species revisited: morphological differentiation among tiger salamanders (*Ambystoma tigrinum*)(Amphibia: Caudata). *Herpetologica* **53**, 30–49 (1997).
31. E. R. Dunn, The races of *Ambystoma tigrinum*. *Copeia* **1940**, 154–162 (1940).
32. E. H. Taylor, Two new ambystomid salamanders from Chihuahua. *Copeia* **3**, 143–146 (1941).
33. R. Webb, Observations on tiger salamanders (*Ambystoma tigrinum* complex, family Ambystomatidae) in Mexico with description of a new species. *Bull. Maryl. Herpetol. Soc.* **40**, 122–143 (2004).
34. P. M. Hime, *et al.*, Phylogenomics reveals ancient gene tree discordance in the amphibian tree of life. *Syst. Biol.* (2020) https:/doi.org/10.1093/sysbio/syaa034 (May 12, 2020).
35. M. Kearse, *et al.*, Geneious Basic: An integrated and extendable desktop software platform for the organization and analysis of sequence data. *Bioinformatics* **28**, 1647–1649 (2012).

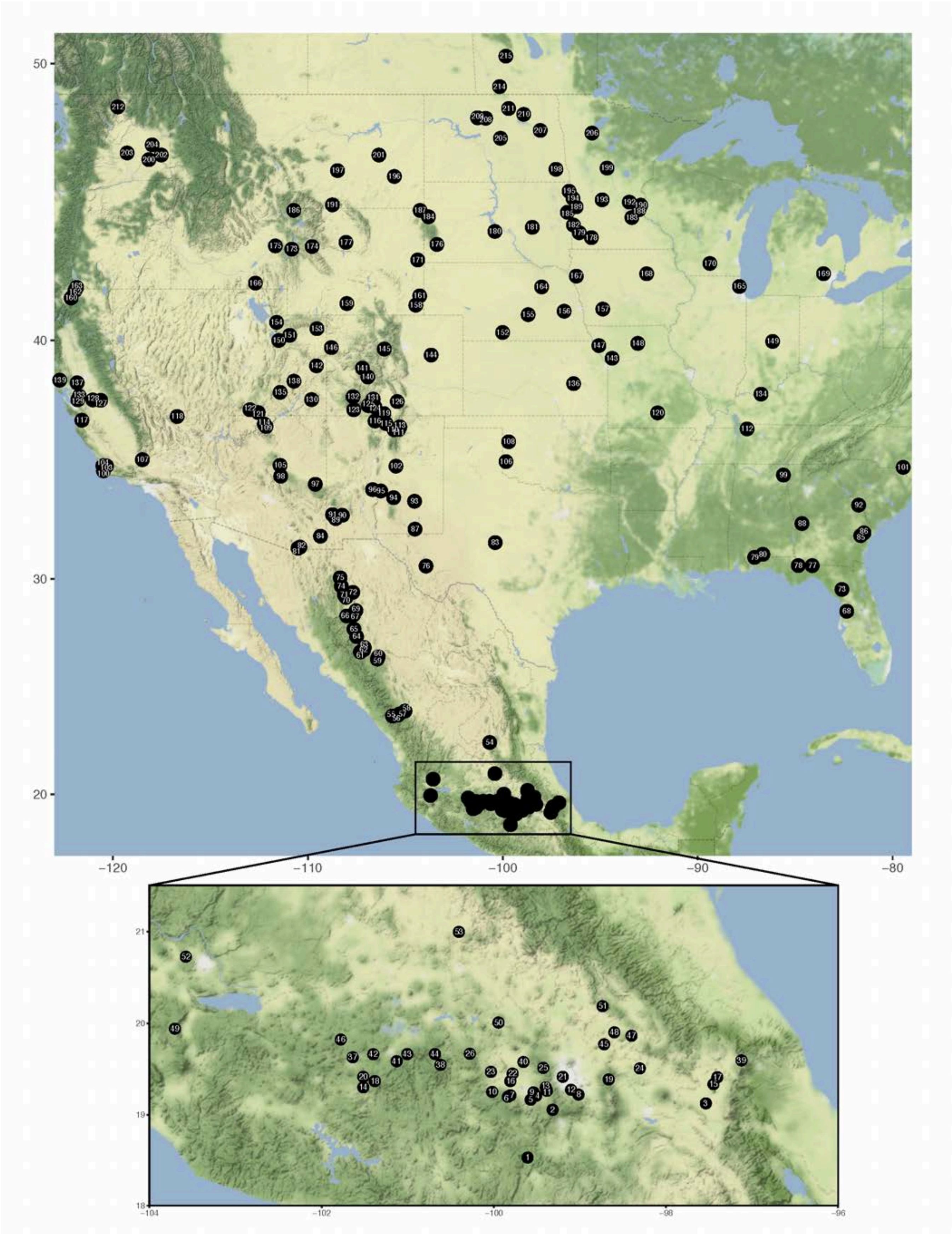


**Figure S1.** Map of collection localities. Numbers correspond to Table S1. The zoomed box gives a detailed view of localities in the Trans-Mexican Volcanic Belt.

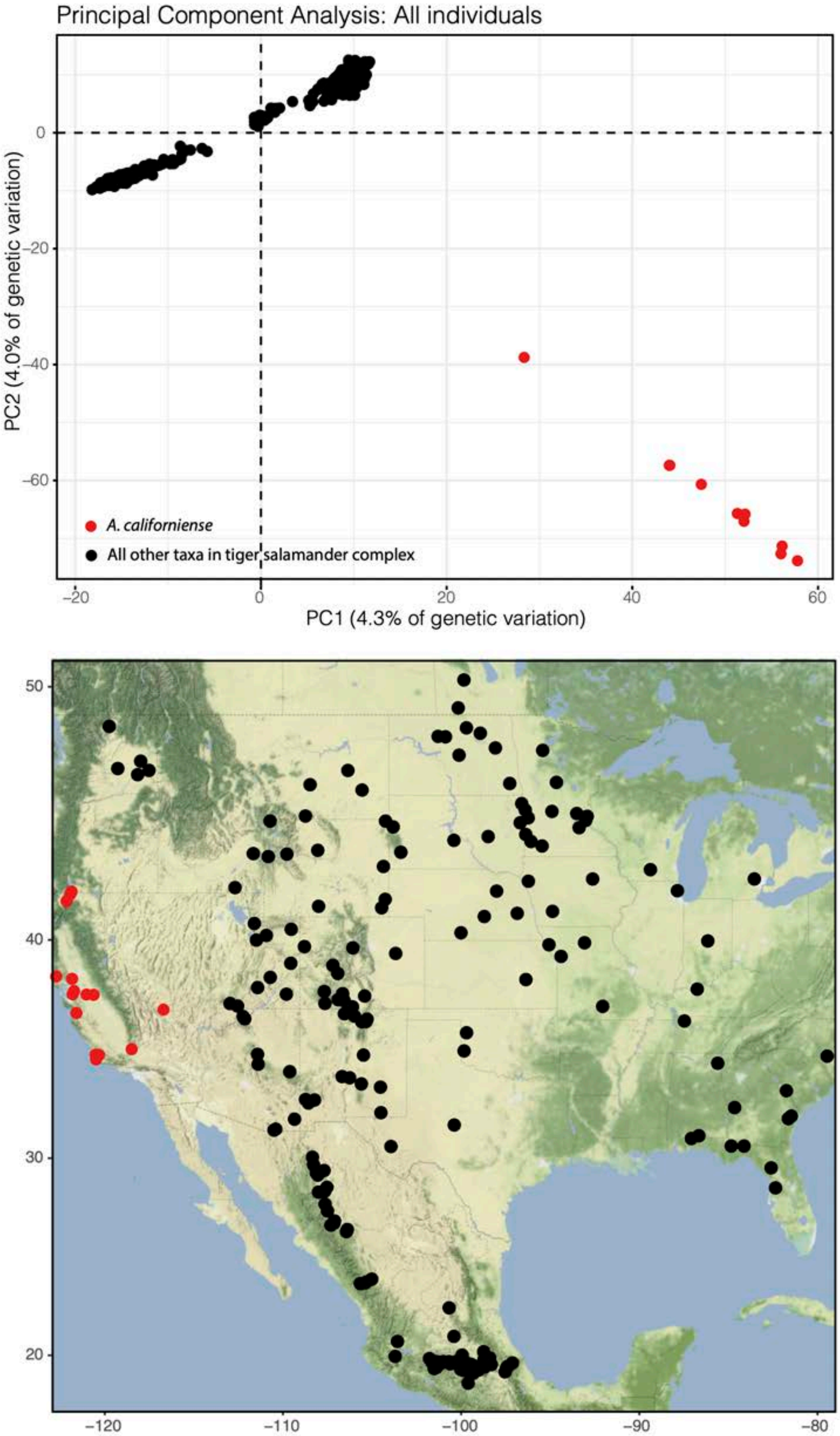


**Figure S2.** Results from the initial principal components (PC) analysis which used the full genetic dataset (all individuals except outgroups). Top graph shows the first (x-axis) and second (y-axis) principal components. Individuals of *A. californiense* are shown as red dots on both the plot and the map below. All other individuals are black.

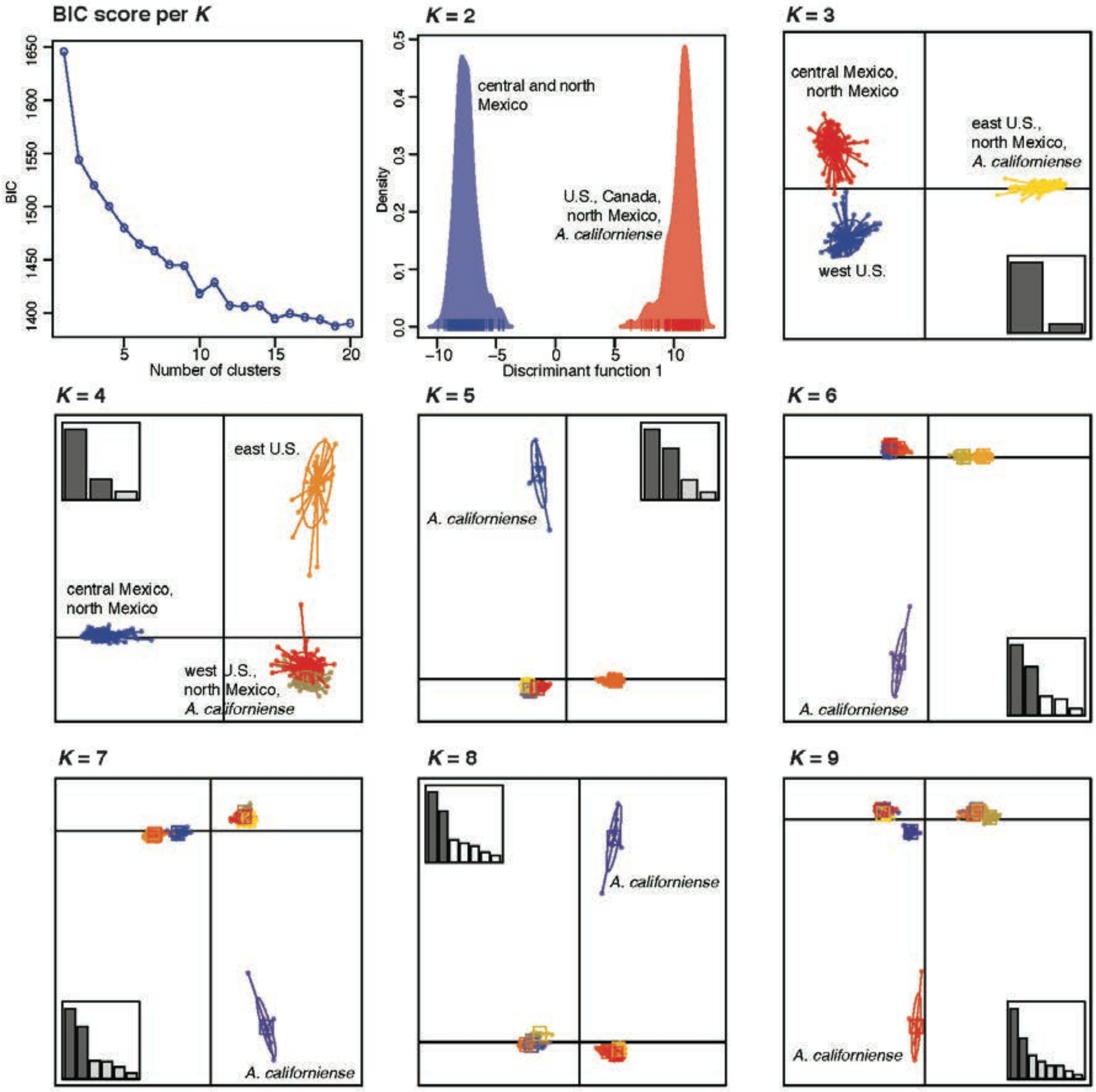


**Figure S3.** DAPC results from exploratory analyses of the full genetic dataset, which included the California tiger salamander (*A. californiense*). The upper left plot shows the BIC scores for each number of possible genetic clusters ($K$). Remaining plots show the first (x-axis) and/or second (y-axis) discriminant functions for various values of $K$. Barplots of discriminant function eigenvalues are inset on each scatterplot. Note that all plots stabilize starting at $K = 5$ in recovering a distinct cluster representing all California tiger salamanders. This trend continued at higher values of $K$.

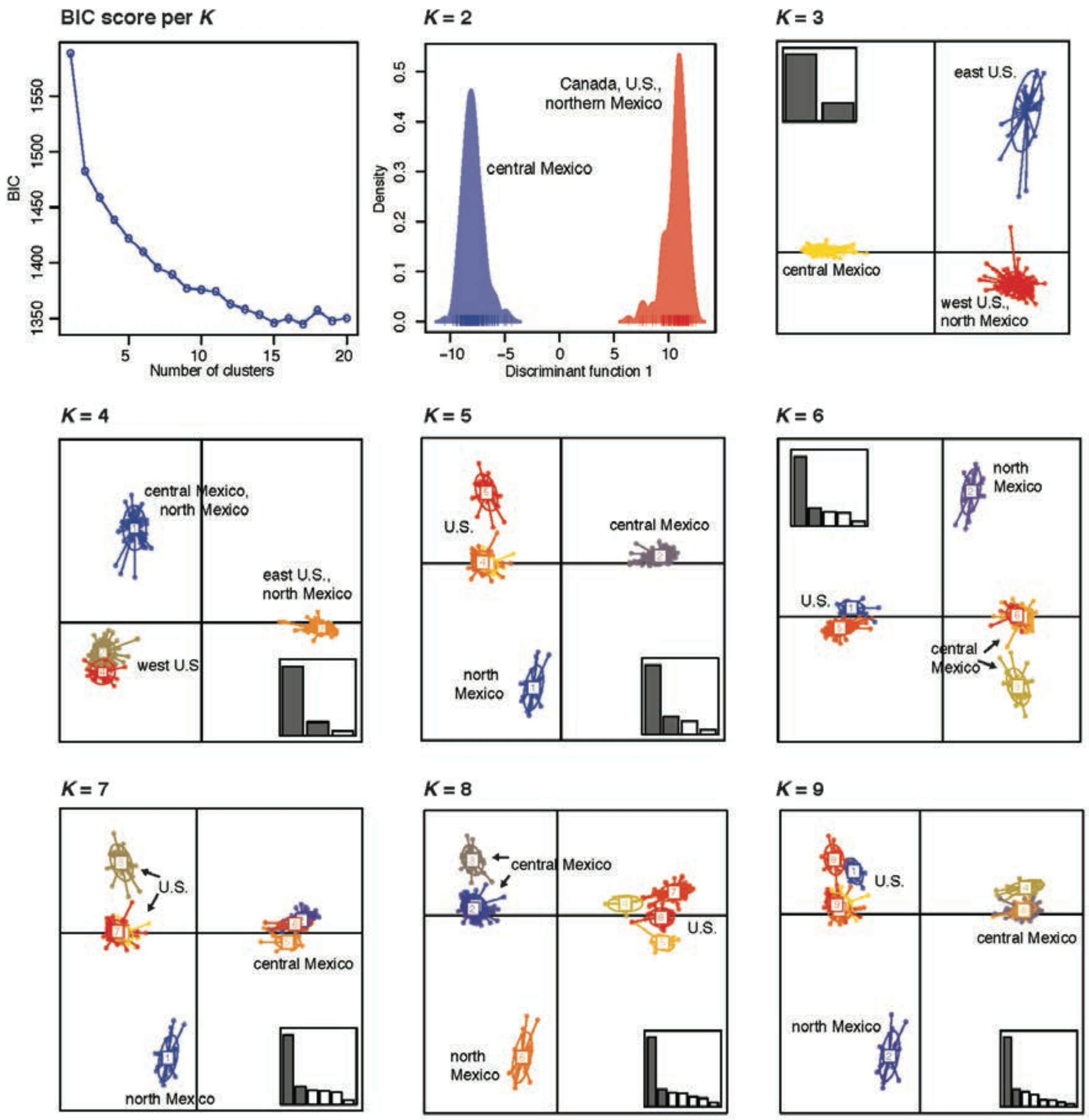


**Figure S4.** DAPC results from exploratory analyses of the tiger salamander complex dataset with *A. californiense* removed. The upper left plot shows the BIC scores for each number of possible genetic clusters (*K*). Remaining plots show the first (x-axis) and/or second (y-axis) discriminant functions for various values of *K*. Barplots of discriminant function eigenvalues are inset on each scatterplot. Note that all plots stabilize starting at *K* = 5 in recovering three predominant genetic clusters representing the U.S. + Canada, northern Mexico, and central Mexico. This trend continued at higher values of *K*.

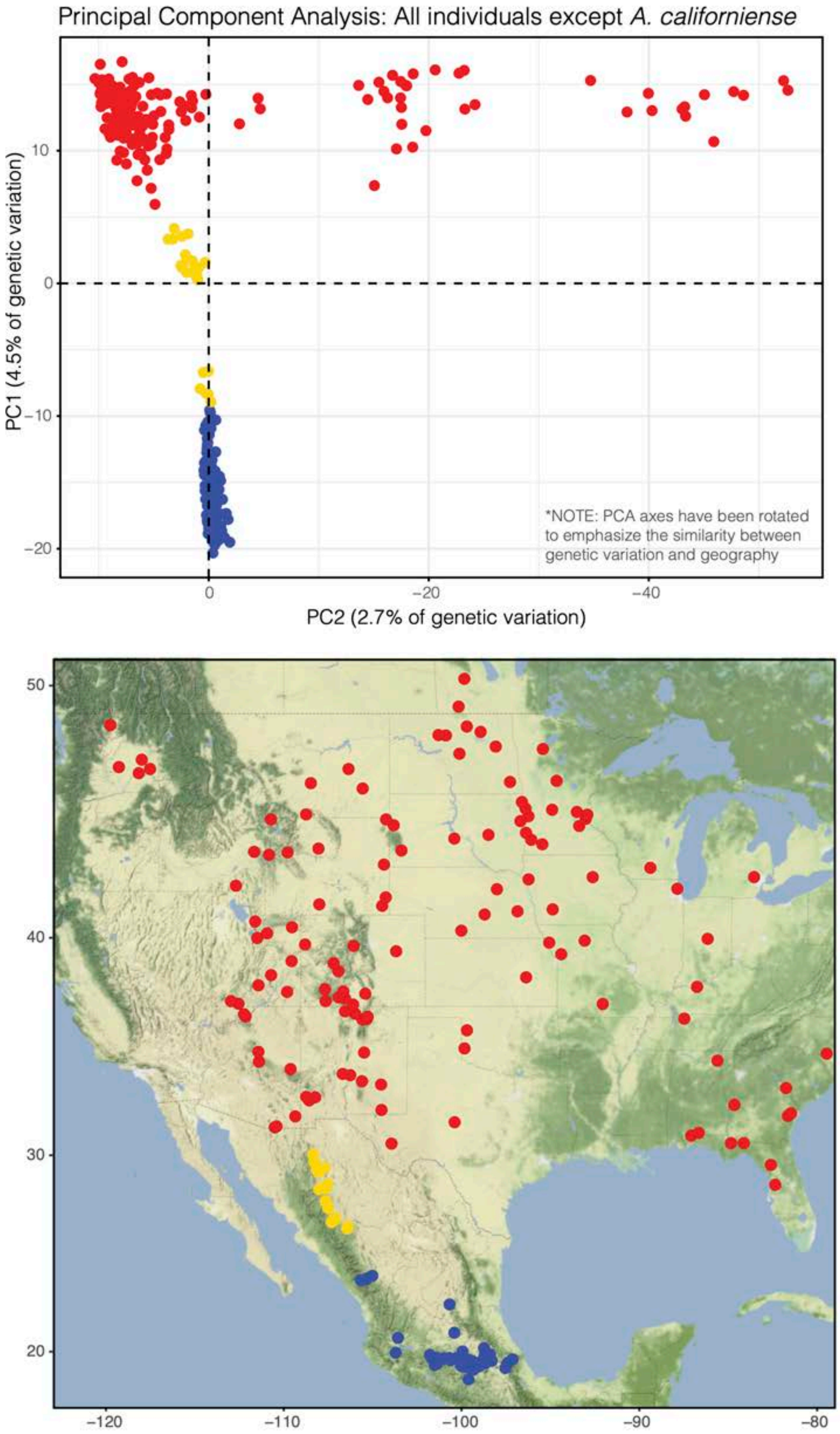


**Figure S5.** Results from the principal components (PC) analysis that included all individuals except outgroups and *A. californiense*. Top graph shows the first (x-axis) and second (y-axis) principal components. Individuals from central Mexico, northern Mexico, and the U.S. + Canada are colored blue, yellow, and red, respectively.

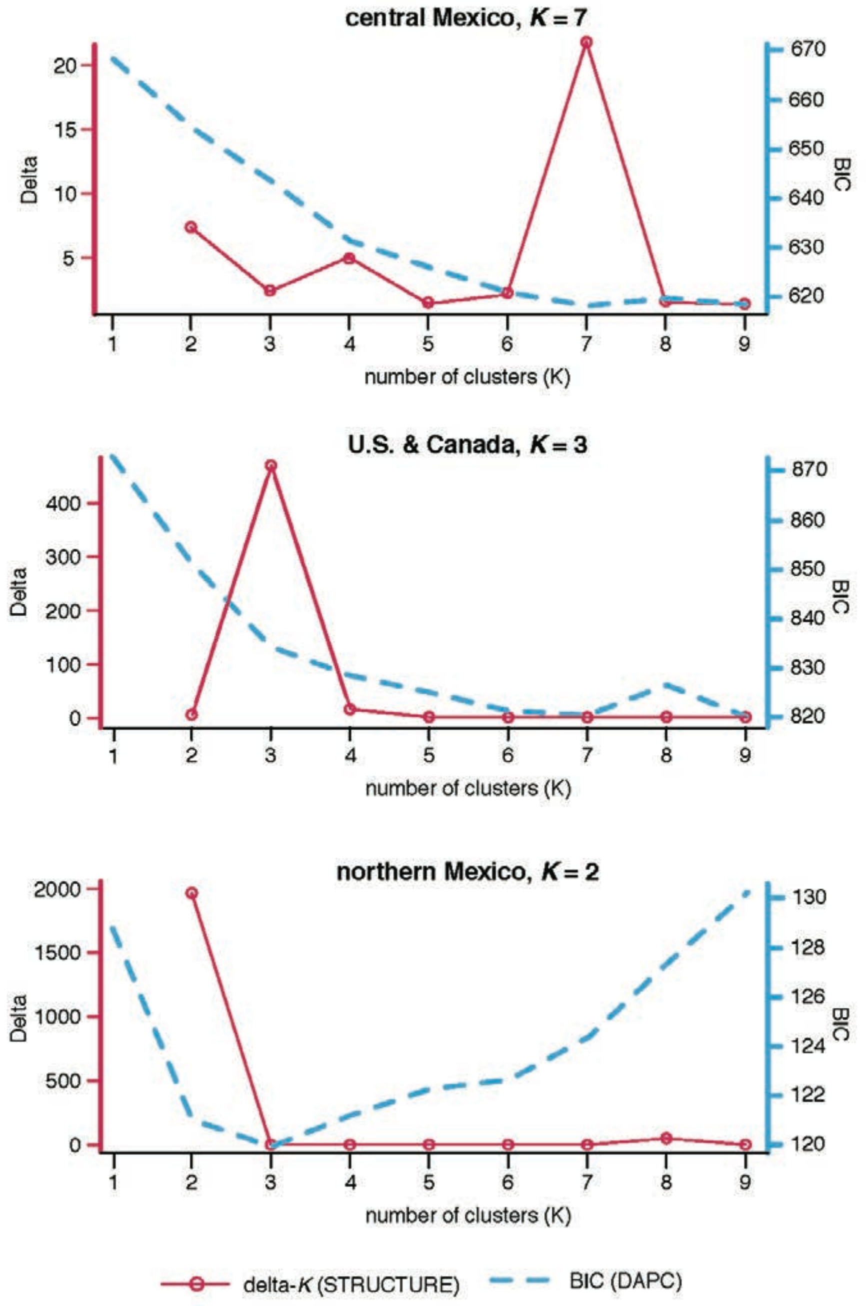


**Figure S6.** Plots used to determine the number of major clusters (*K*) within the central Mexico, U.S., and north Mexico groups (top, middle, and bottom rows, respectively). The left column contains plots of delta-*K*, calculated using the method of Evanno et al. (2005), used to determine the number of clusters in STRUCTURE analyses. The right column contains plots of BIC, used to determine the number of clusters in DAPC analyses. Membership plots and maps relating to these results are shown in Fig. 1.

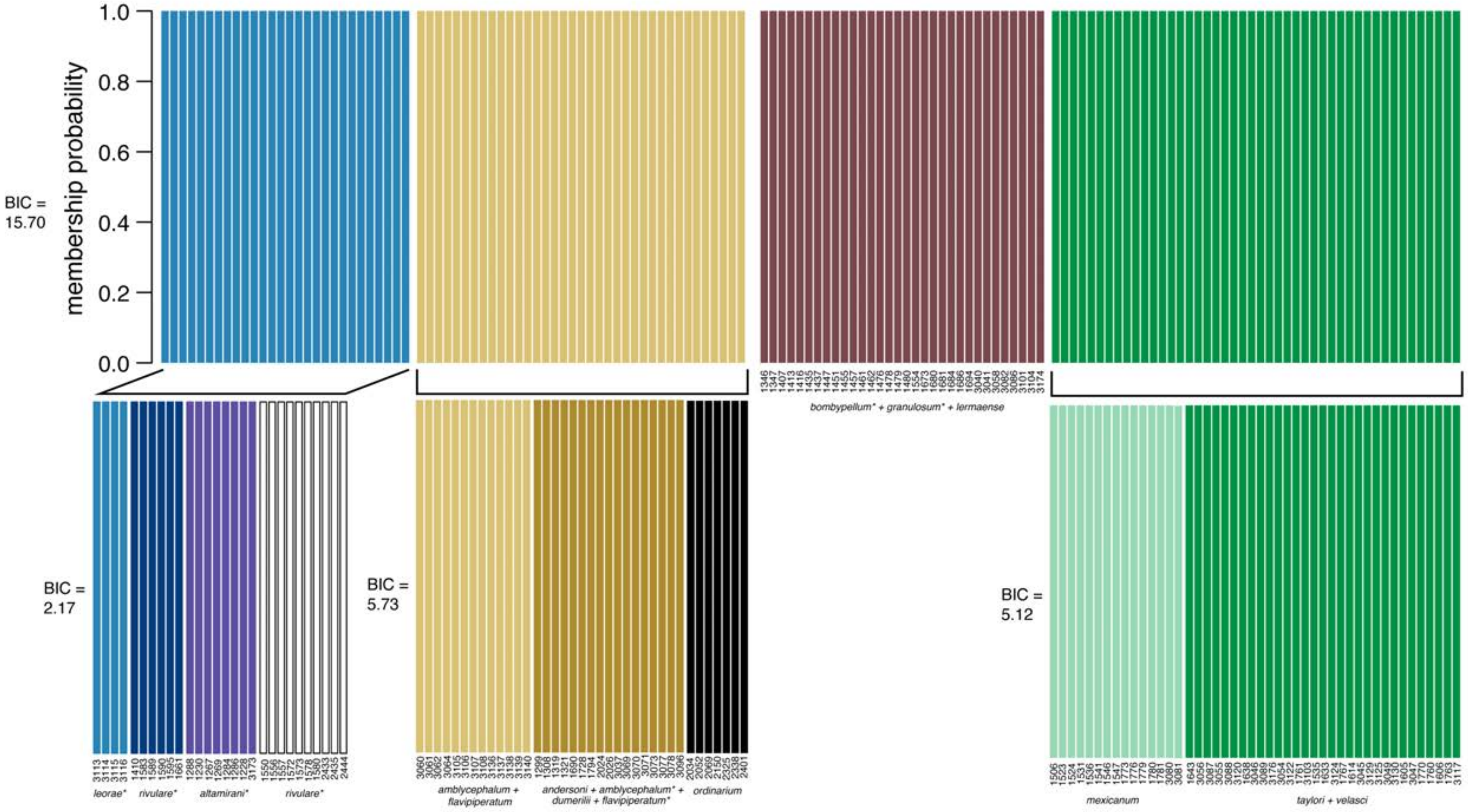


**Figure S7.** Results of discriminant analysis of principal components (DAPC) for the Central Mexico genetic subgroup. Recursive rounds of DAPC analyses (indicated by brackets) were performed as described in the main text. Values of delta-BIC for each round are given to the left of each membership plot. Note that no further rounds of analyses were conducted after the value of delta-BIC fell below 2. Taxonomic names listed below some clusters are based on the presence of individuals collected from (or near, indicated by *) the type locality of that species or subspecies.

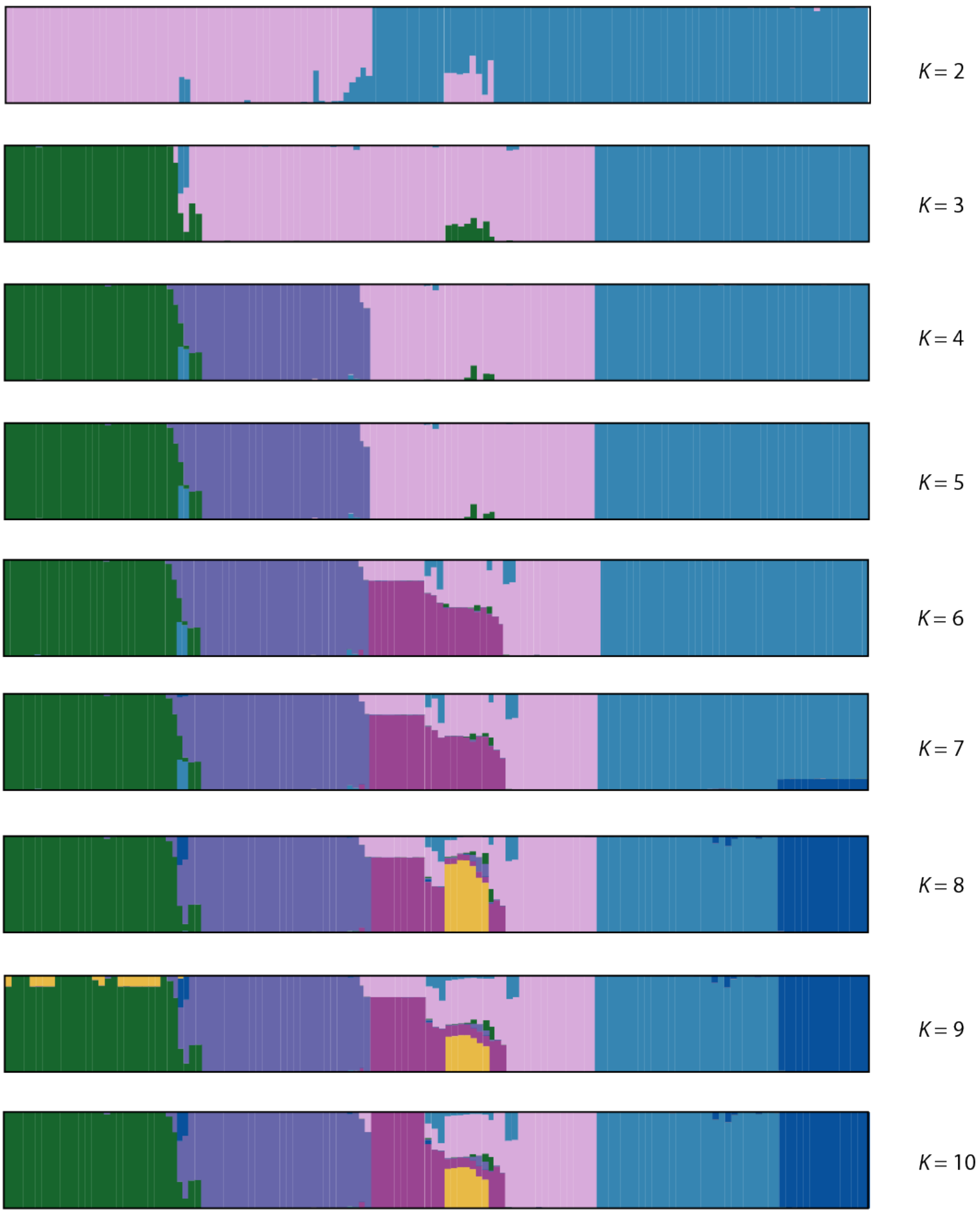


**Figure S8.** Results from exploratory STRUCTURE analyses of the central Mexico, from *K* = 1 to *K* = 10. Each vertical bar represents an individual, while the y-axis gives the probability of group membership. The best-supported number of clusters was *K* = 7 (Fig. S6), however, a pattern of four primary clusters emerged starting at *K* = 4; these were the four clusters used in further analyses.

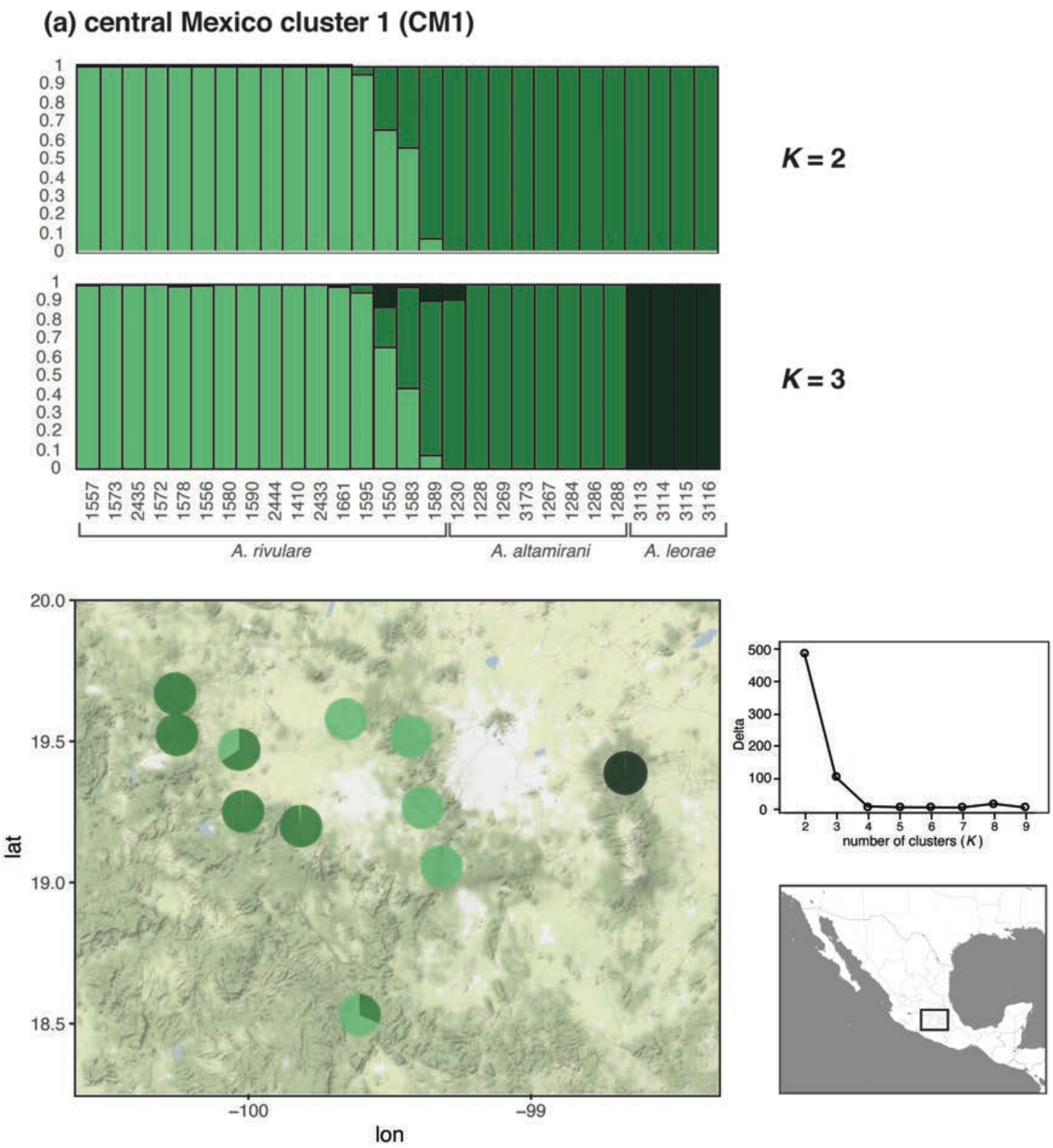


**Figure S9.** Detailed results from exploratory follow-up STRUCTURE analyses of (a) CM1, (b) CM2, (c) CM3, and (d) CM4 (see Fig. 2 for results from prior rounds of analyses). Each vertical bar represents an individual, while the y-axis gives the probability of group membership. Lower values of *K* were selected using *a priori* selection methods described in the text, while higher values of *K* represent the number of currently described species in that cluster. Species names below membership plots represent the taxonomic designations that would have been made in the field. Plots of delta(*K*) calculated using the method of Evanno et al. (2005) are to the right of each map.

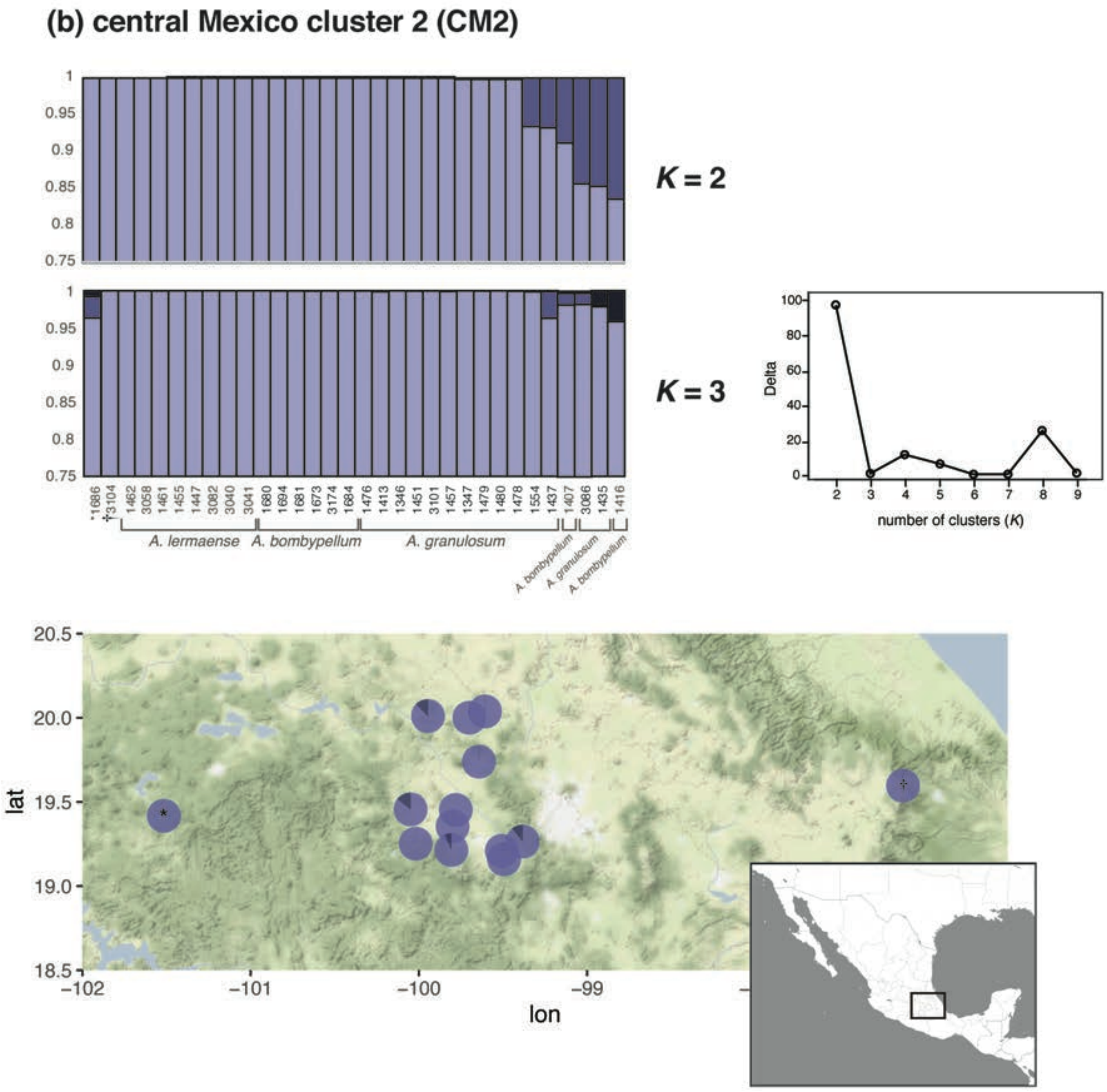


**Figure S9 (continued).** Individuals 1686 (*) and 3104(†) are range outliers that were not shown in Fig. 1 (see discussion in supplementary text).

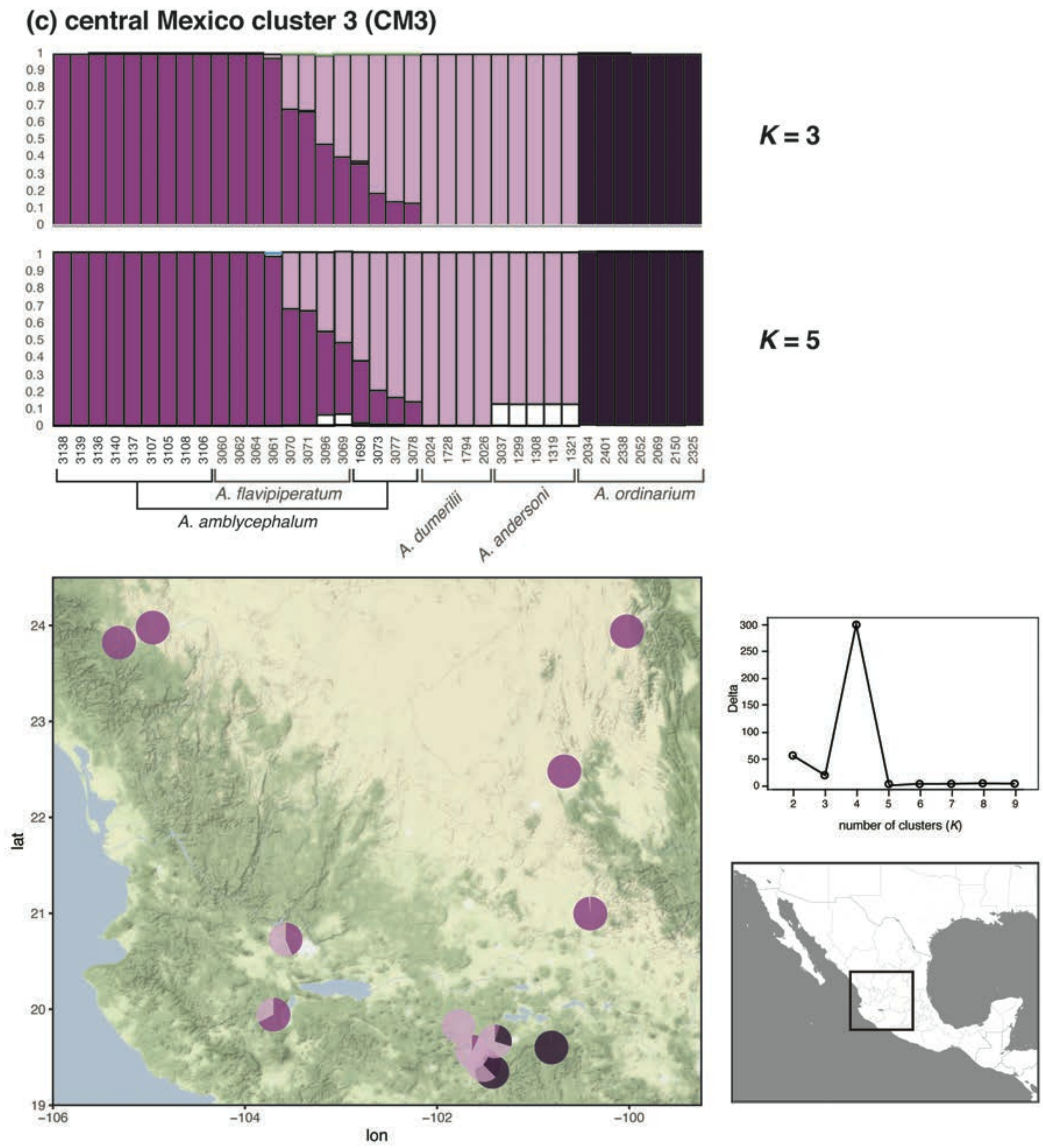


392

393 **Figure S9 (continued).**

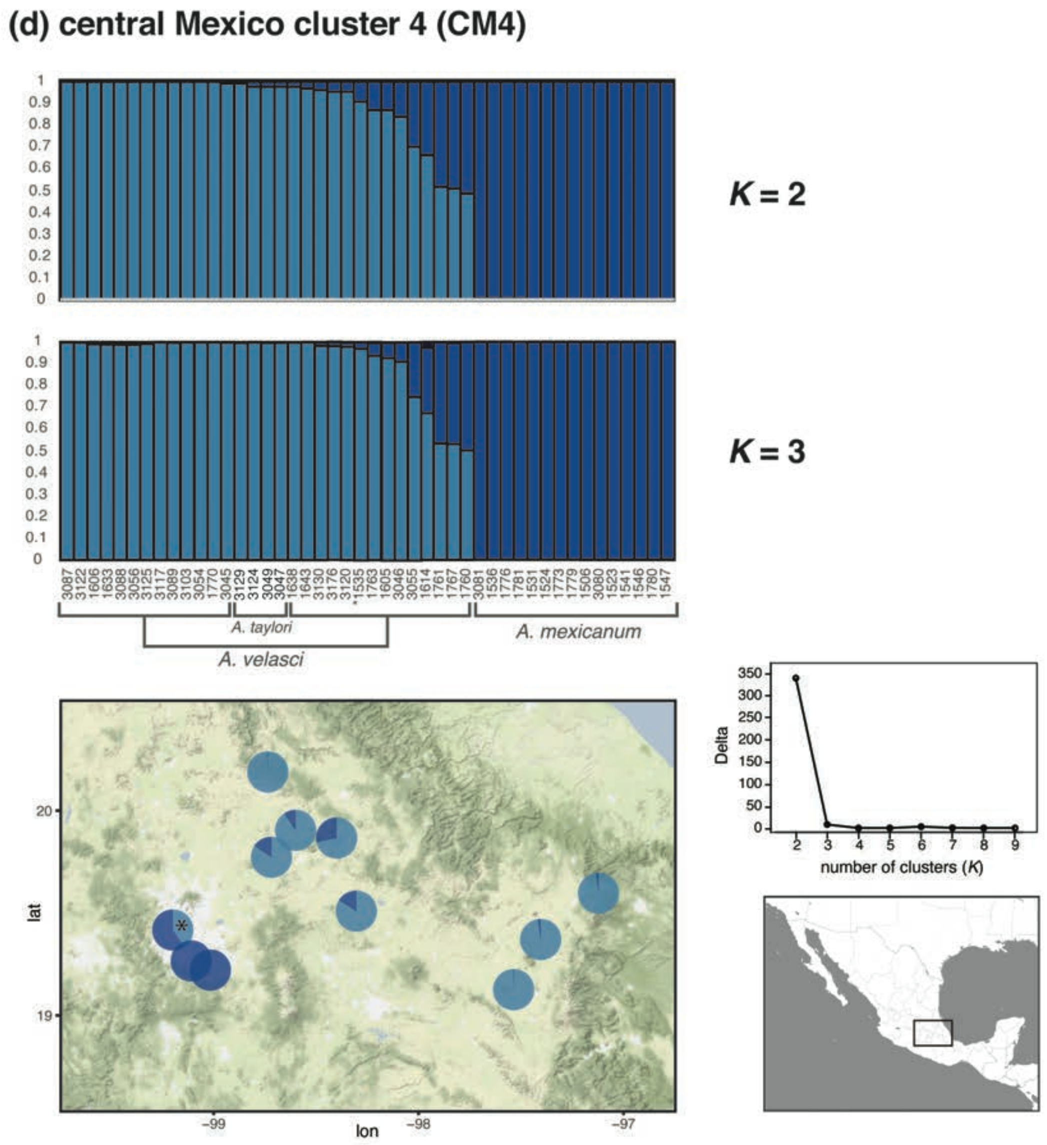


**Figure S9 (continued).** Individual 1535(†) was collected from Chapultepec (a locality where only *A. mexicanum* was expected to occur) yet falls out in the major genetic cluster that includes *A. taylori* (lighter blue on the top STRUCTURE membership plot).

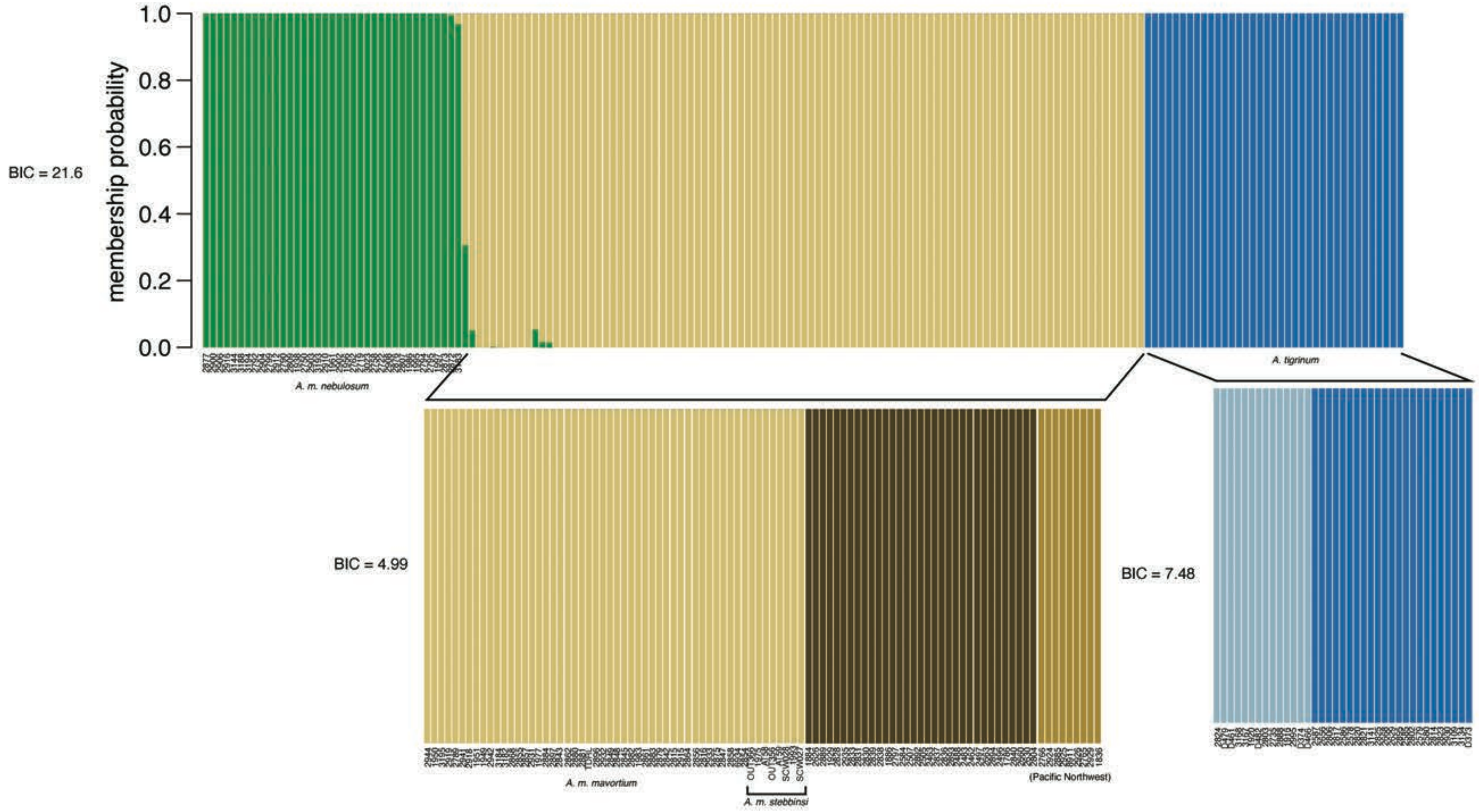


**Figure S10.** Results of discriminant analysis of principal components (DAPC) for the U.S. genetic subgroup. Recursive rounds of DAPC analyses (indicated by brackets) were performed as described in the main text. Values of delta-BIC for each round are given to the left of each membership plot. Note that no further rounds of analyses were conducted after the value of delta-BIC fell below 2. Taxonomic names listed below some clusters are based on the presence of individuals collected from the type locality of that species or subspecies.

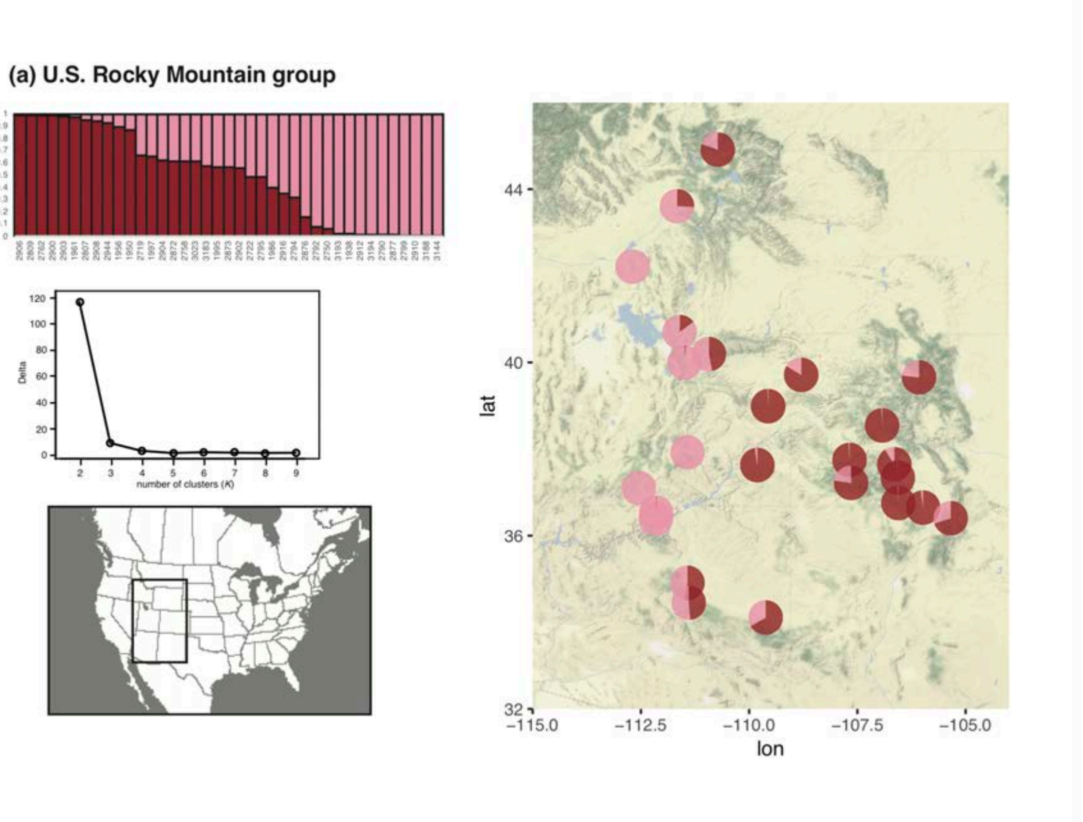


**Figure S11.** Detailed results from exploratory follow-up STRUCTURE analyses of (a) U.S. Rocky Mountain group, (b) U.S. central group, and (c) U.S. eastern group (see Fig. 2 for results from the first round of analyses). Each vertical bar represents an individual, while the y-axis gives the probability of group membership. Plots of delta(*K*), calculated using the method of Evanno et al. (2005), are also provided below each membership plot.

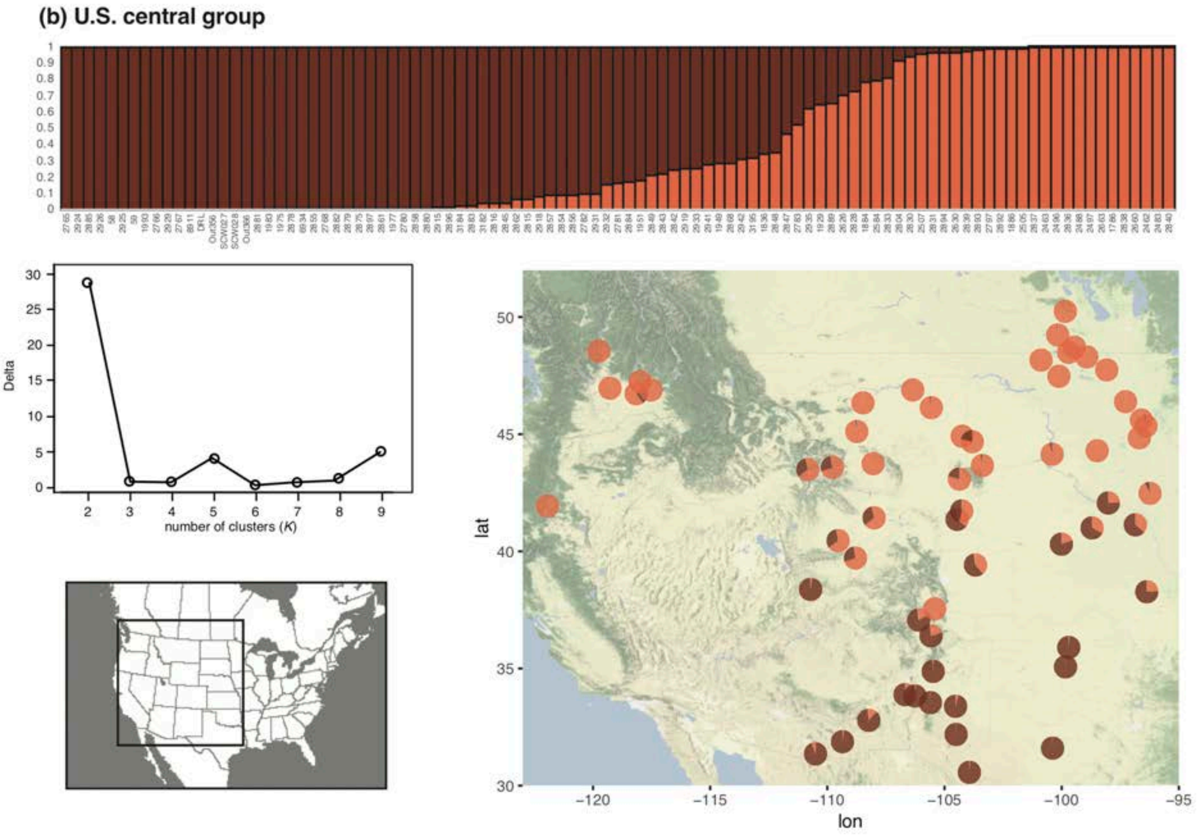


**Figure S11 (continued).**

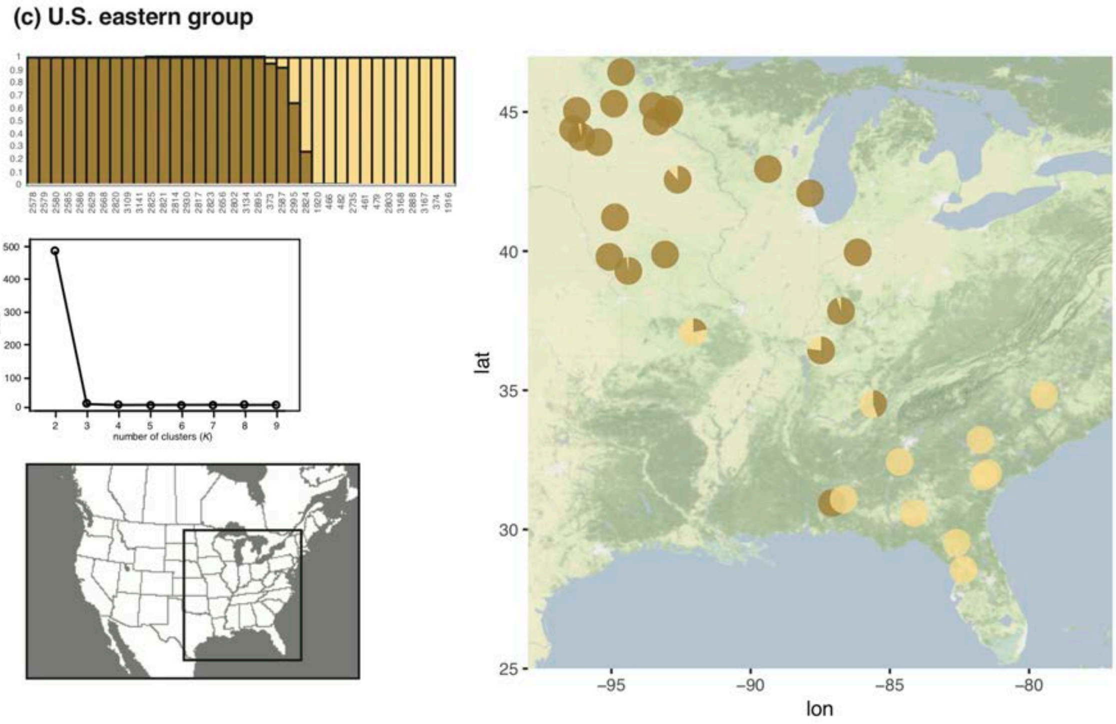


**Figure S11 (continued).**

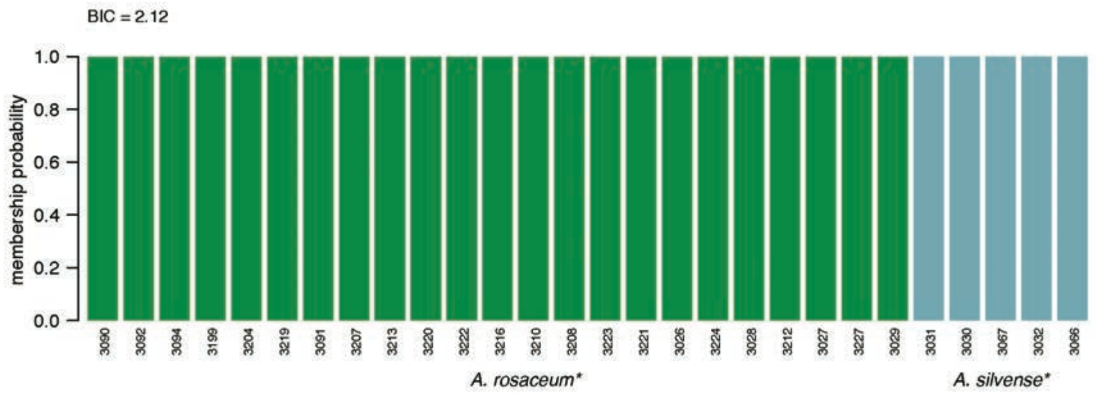


**Figure S12.** Results of discriminant analysis of principal components (DAPC) for the northern Mexico genetic subgroup. Value of delta-BIC is given above the membership plot. Note that no further rounds of analyses were conducted after the value of delta-BIC fell below 2. Taxonomic names listed below clusters are based on the presence of individuals collected from near (indicated by *) the type locality of that species or subspecies.

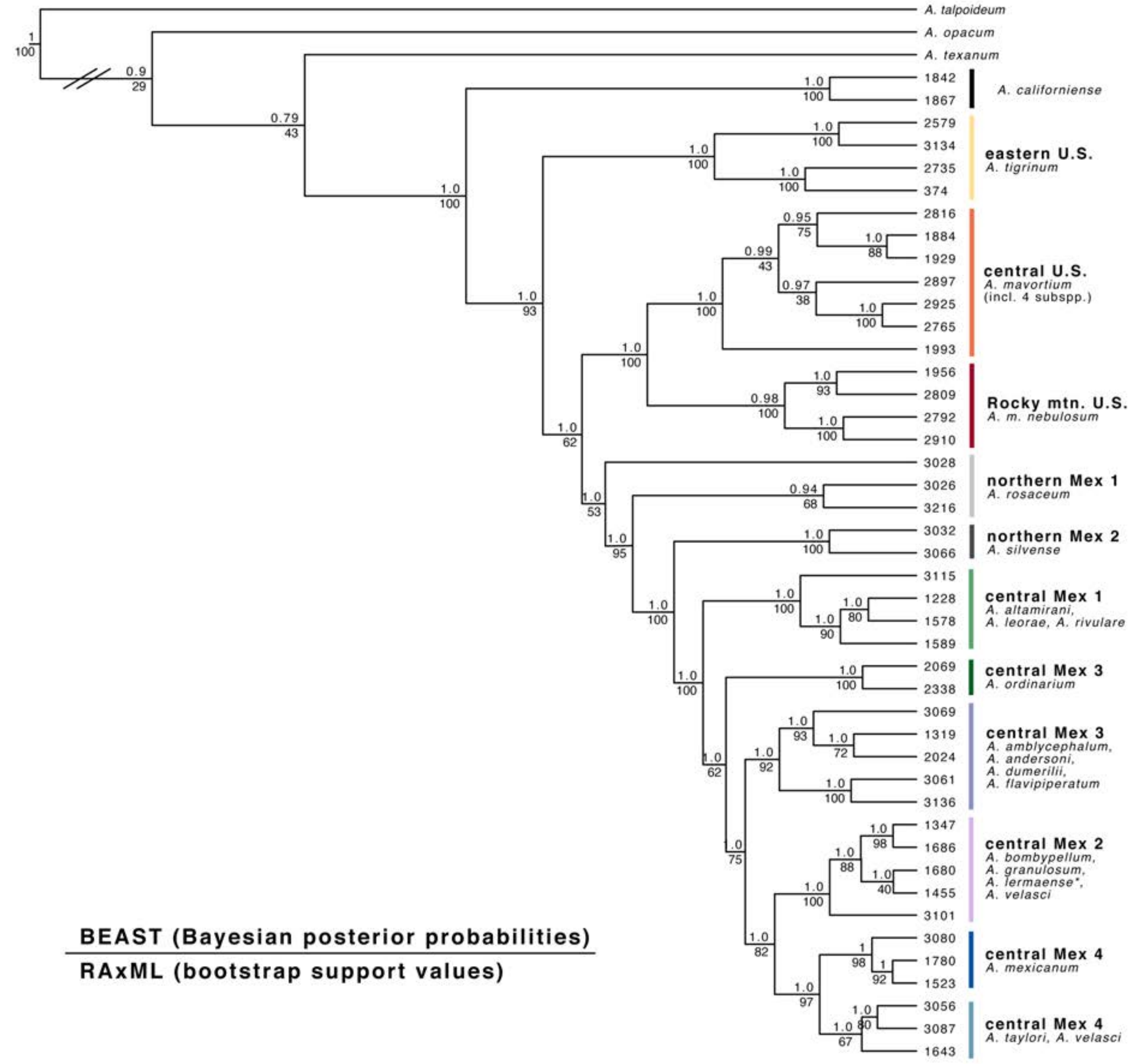


**Figure S13.** Results of concatenated phylogenetic analyses performed using the Bayesian program BEAST and the maximum-likelihood program RAxML. Both methods produced identical topologies. Bayesian posterior probabilities and bootstrap support values are provided above and below each node, respectively. Branch lengths were produced by BEAST. Colored clades correspond to the major genetic clusters (and associated species) identified by population genetic analyses (Fig. 2).

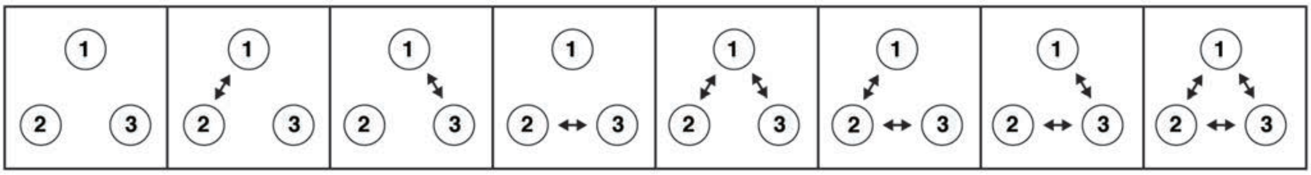

**Figure S14.** A graphical representation of the models evaluated in Migrate-N analyses. Models vary in the presence or absence of gene flow among populations, and model fit was assessed using Bayes Factors (Table S3). In the analysis of CM3, population 1 is *Ambystoma andersoni* and population 2 is *A. dumerilii*. In the analysis of CM4, population 1 is *A. mexicanum* and population 2 is *A. taylori*. In both analyses, population 3 represents facultatively paedomorphic taxa from surrounding localities in the same genetic cluster.

**Table S1.** Lab IDs (DWW#), field numbers, taxonomic assignments, and locality information for specimens used in this study. This table is available on Figshare (https://figshare.com/s/abb195a2464c55ddcc40).

**Table S2.** The number of filtered sequencing reads associated with each individual/locus combination. This table is available on Figshare (https://figshare.com/s/9f3dd2cec2600234ca94).

**Table S3.** Natural logarithms of Bézier-corrected marginal likelihoods (Bezier lnL), Bayes factors (BF), and model ranks for each of the models evaluated in migrate-n (described in text and Fig. S14). The top-ranked models are shaded gray.

| | CM3 | | | CM4 | | |
|---|---|---|---|---|---|---|
| **Model Number** | **Bezier lnL** | **BF** | **Rank** | **Bezier lnL** | **BF** | **Rank** |
| Model 1 *(no migration)* | -40759.29 | -1461 | 8 | -42348.67 | -3711 | 8 |
| Model 2 | -40639.73 | -1222 | 7 | -41978.36 | -2971 | 7 |
| Model 3 | -40376.65 | -696 | 6 | -41388.60 | -1791 | 6 |
| Model 4 | -40276.49 | -495 | 5 | -41222.09 | -1458 | 5 |
| Model 5 | -40057.11 | -57 | 3 | -40511.74 | -37 | 2 |
| Model 6 | -40086.80 | -116 | 4 | -40591.43 | -197 | 3 |
| Model 7 | -40056.16 | -55 | 2 | -40618.06 | -250 | 4 |
| Model 8 (full migration) | -40028.66 | 0 | 1 | -40493.02 | 0 | 1 |

**Table S4.** Parameter estimates from migrate-n analyses for population size ($\Theta$) and mutation-scaled migration rate (*M*). Confidence interval = 95% CI.

| **CM3** | **Median** | **95% CI** |
|---|---|---|
| $\Theta_1$ (*A. andersoni*) | 0.0023 | 0.0002 – 0.0038 |
| $\Theta_2$ (*A. dumerilii*) | 0.0031 | 0.001 – 0.0046 |
| $\Theta_3$ (facultative taxa) | 0.0041 | 0.002 – 0.0056 |
| $M_{1>2}$ | 751 | 640 – 854 |
| $M_{1>3}$ | 441 | 338 – 538 |
| $M_{2>1}$ | 457 | 344 – 578 |
| $M_{2>3}$ | 381 | 290 – 474 |
| $M_{3>1}$ | 481 | 378 – 586 |
| $M_{3>2}$ | 613 | 514 – 718 |

| **CM4** | **Median** | **95% CI** |
|---|---|---|
| $\Theta_1$ (*A. mexicanum*) | 0.0023 | 0.0002 – 0.0038 |
| $\Theta_2$ (*A. taylori*) | 0.0033 | 0.001 – 0.0046 |
| $\Theta_3$ (facultative taxa) | 0.0046 | 0.0022 – 0.0066 |
| $M_{1>2}$ | 841 | 726 – 942 |
| $M_{1>3}$ | 891 | 786 – 978 |
| $M_{2>1}$ | 217 | 148 – 292 |
| $M_{2>3}$ | 679 | 574 – 782 |
| $M_{3>1}$ | 209 | 142 – 274 |
| $M_{3>2}$ | 609 | 496 – 718 |
